# Live-cell imaging provides direct evidence for a threshold in CDK activity at the G2/M transition

**DOI:** 10.1101/2023.03.26.534249

**Authors:** Hironori Sugiyama, Yuhei Goto, Yohei Kondo, Damien Coudreuse, Kazuhiro Aoki

**Author notes:** These authors contributed equally to this work (Hironori Sugiyama, Yuhei Goto).

## Abstract

Cyclin-dependent kinase (CDK) plays an essential role in determining the temporal ordering of the cell cycle phases. However, despite significant progress in studying regulators of CDK, it remains elusive how they coordinately affect CDK activity at the single-cell level and how CDK controls the temporal order of cell cycle events. This could be due to the lack of tools to monitor CDK activity in living cells. Here, we elucidate the dynamics of CDK activity in fission yeast and mammalian cells by using a newly developed CDK activity biosensor, Eevee-spCDK, based on Förster Resonance Energy Transfer (FRET). Taking advantage of this system, we unravel the profile of CDK activity in vegetatively growing *S. pombe* cells. Thus, we detect a transient increase in S phase followed by a gradual increment during G2 phase. CDK activity then reaches its maximum in early M phase and rapidly decreases at mitotic exit. During G2 phase, CDK activity exhibits a biphasic pattern, *i.e.*, an early slow increase and a late fast rise prior to the G2/M phase transition, as predicted from mathematical studies. Remarkably, although CDK activity does not necessarily correlate with cyclin levels, we find that it converges to the same level around mitotic onset in several mutant backgrounds, including *pom1Δ* cells and *wee1* or *cdc25* overexpressing cells. These data provide the first direct evidence that cells enter M phase when CDK activity reaches a high threshold, consistent with the quantitative model of cell cycle progression in fission yeast.

## Introduction

The cell cycle is a highly conserved biological process that controls cell proliferation, and proper regulation of cell cycle progression is critical to maintain homeostasis in cell size and other physiological features^1^. In eukaryotes, cyclin-dependent kinases (CDK) play an essential role in cell cycle control^2^. Thus, cells sense environmental cues, such as nutrients and growth factors as well as stress molecules, and integrate the information through modulating CDK activities to determine whether they will divide or not. CDKs are primarily regulated by direct association with members of the cyclin family, which are required for CDK catalytic activity, thereby driving progression through the cell cycle via the phosphorylation of key CDK substrates^2, 3^. Different types of cyclins accumulate and associate with different types of CDKs at each cell cycle phase^3^. In addition to cyclins, other CDK regulators impact cell cycle progression by post-translational modification or direct binding to CDKs^4^.

The fission yeast *Schizosaccharomyces pombe* (*S. pombe*) is a particularly suitable model organism for eukaryotic cell cycle research. This is mainly because *S. pombe* contains only one CDK (encoded by *cdc2*) to drive the cell cycle^5, 6^. Furthermore, among the four main cell cycle cyclins (Cdc13, Puc1, Cig1, and Cig2), only the B-type cyclin Cdc13 is essential for the *S. pombe* cell cycle^7–11^. Given the conservation of the core cell cycle regulation among eukaryotes^12^, the simplicity of the fission yeast cell cycle network is a unique asset to decipher the essence of cell cycle mechanisms^3^. In part for this reason, the question of how fission yeast cells regulate their division cycle has long attracted attention in cell biology. Interestingly, despite the diversity in cyclin-CDK pairs at specific stages of the cell cycle, it has been proposed that different levels of CDK activity control fission yeast cell cycle progression, with a low-to-moderate level of CDK activity initiating S phase entry (G1/S transition), and a further increase in CDK activity to a high level triggering mitosis^13^ (G2/M transition). As mentioned above, in addition to cyclin binding, CDK activity is modulated by post-translational modification of CDK. Thus, CDK function during G2 and M phases is altered by a conserved network of two feedback loops that control the phosphorylation state of the Tyr15 (Y15) residue of CDK^14^: the kinase Wee1 inactivates CDK by phosphorylating Y15 while the phosphatase Cdc25 activates CDK through dephosphorylation of Y15^15^.

These two regulatory mechanisms, namely cyclin binding and post-translational modifications of CDKs, have been separately reported to be involved in determining the timing of cell division. Notably, it was proposed that cyclin concentration correlates with cell size, and that cell division occurs above a certain cyclin concentration threshold^16^. Furthermore, the mitotic inhibitor Pom1, a member of the DYRK family kinases, was suggested to indirectly modulate Wee1 activity, contributing to CDK regulation and cell size control through CDK post-translational modification^17, 18^. However, the exact mechanism is still under debates^19–21^. In addition, the gradual accumulation of Cdc25 during G2 phase, which counteracts the Wee1 function, plays a role in determining the timing of mitosis^22, 23^. Despite these extensive studies of such key regulators of CDK, it remains elusive how CDK controls the temporal order of cell cycle events because of the combinatorial complexity of CDK regulations. The involvement of phosphatases, such as PP1 and PP2A^24^, makes the situation even more complicated, as the overall phosphorylation output of the CDK system is determined by a balance between the activity of CDKs and that of CDK-counteracting phosphatases. Importantly, the dynamic behavior of CDK activity, integrating the effects of its regulators and those of conserved phosphatases, has rarely been measured at the single-cell level, mainly due to a lack of tools to directly quantify CDK activity in individual living cells.

Genetically encoded biosensors for CDK have been developed, mainly in more complex eukaryotes^25^, enabling the *in vivo* visualization of CDK activity. First, CDK1 activity in mammalian cells was detected by biosensors based on the principle of Förster Resonance Energy Transfer (FRET) ^26, 27^. Another FRET biosensor for CDK1 was also recently applied in a reconstituted system with *Xenopus* egg extracts^28^. In cultured mammalian cells, translocation-based biosensors allowed for monitoring CDK1/2^29, 30^ and CDK4/6^31^ activities. Surprisingly, while the simplicity of the fission yeast cell cycle regulatory network and the host of tools for advanced genetic engineering available in *S. pombe* allow for in-depth investigation of the core principles underlying cell cycle progression in eukaryotes, FRET biosensor for CDK were never established for this organism. A translocation-based biosensor, synCut3, is the only tool that was developed thus far to prove CDK activity at the single-cell level in fission yeast^32^. Importantly, synCut3 mainly detects CDK activity during mitosis, showing an increase just prior to mitotic entry and a decrease at mitotic exit. However, CDK plays an important role in cell cycle progression not only at mitotic entry, but also in the S and G2 phases. In fact, the complete fission yeast cell cycle can be executed by a single cyclin-CDK complex, suggesting that quantitative differences in CDK activity define independent cell cycle phases^11^. Furthermore, it has been demonstrated that the CDK activity threshold quantitatively differs among CDK substrates, and such differences in activity threshold may contribute to the temporal ordering of cell cycle phases^33^. Therefore, to understand how CDK governs cell cycle progression, CDK biosensors for *S. pombe* must be developed to monitor the quantitative behavior of CDK activity throughout the cell cycle phases at the single-cell level.

Here, we report a newly developed FRET biosensor for CDK, named Eevee-spCDK, which allowed us to visualize CDK activity in fission yeast. FRET imaging with Eevee-spCDK demonstrated that in vegetatively growing *S. pombe* cells, CDK activity—or the balance between CDK activity *per se* and related phosphatases— increases at the time of S phase, then gradually rises during G2 to reach a maximum level in M phase. Eevee-spCDK further revealed that the increase in CDK activity during G2 is biphasic. Remarkably, we found that this pattern of CDK activity during the cell cycle is robust even in conditions where cell size is perturbed or upon overexpression of Cdc25/Wee1, while cyclin concentration was uncoupled from cell size and even CDK activity. Eevee-spCDK was further shown to allow for monitoring CDK1 and CDK2 activity in mammalian cells.

## Results

### Development of a CDK biosensor in *S. pombe*, Eevee-spCDK

To quantitatively monitor the dynamics of CDK activity during the cell cycle, we set out to construct a FRET biosensor for CDK in fission yeast (Figure 1A). In brief, the FRET biosensor contains YPet, a yellow fluorescent protein variant (YFP), and Turquoise2-GL, a cyan fluorescence protein (CFP) variant, as an acceptor and a donor, respectively. These fluorophores are separated by a phosphoserine– proline or phosphothreonine–proline binding domain (WW domain from human Pin1^34^), a long flexible linker (EV linker^35^), and a substrate domain phosphorylated by CDK (sensor domain). In addition, we use a nuclear localization signal (NLS) at the C-terminal of the biosensor, as CDK is predominantly localized in the nucleus throughout the cell cycle in *S. pombe*^36^. When CDK phosphorylates the sensor domain, the WW domain recognizes the phosphorylated substrate and brings CFP in close proximity to YFP, leading to energy transfer from CFP to YFP. Therefore, this type of biosensor monitors the balance between the phosphorylation activity of CDK and the activity of its counteracting phosphatase(s).

First, we investigated what sensor domain may be the most suitable for a CDK FRET biosensor. Among various CDK substrates characterized by previous phospho-proteomic studies^33^, we selected 7 candidate CDK substrates (Drc1, Rad2, Orc1, Orc2, Cut3, Alp14, and Ask1). 12 peptides from these putative substrates were used to build an array of potential CDK biosensors. Each peptide consists of approximately 20 amino acids and includes the phosphoacceptor serine or threonine residue (Figure 1B). Previous reports have shown that CDK activity is correlated with cyclin levels^16, 32^. Therefore, to screen for the response of the different FRET biosensors to CDK activity, we examined the correlation between the FRET/CFP ratio (see Materials and Methods), a good proxy for FRET efficiency, and the amount of endogenous cyclin (*i.e.*, Cdc13-mScarlet-I). However, none of the constructed FRET biosensors gave signals that correlated with cyclin levels (Figure 1C), implying that a single CDK phosphorylation site may not be sufficient to generate a detectable change in FRET efficiency. We therefore tested another set of constructs integrating domains from Drc1 (Drc1N) and Cut3 (Cut3N) as sensor domains, as these contain multiple CDK target sites (Figure 1D). Remarkably, the biosensor containing Drc1N showed an increase in FRET/CFP ratio as the cyclin level increased, with a gain of approximately 40%. In contrast, the Cut3N-containing biosensor did not display any change in FRET/CFP ratio (Figure 1E). We named the Drc1N-containing biosensor Eevee-spCDK, as it is based on the Eevee backbone for intramolecular FRET biosensors^35^. Live-cell imaging revealed a positive correlation between the FRET/CFP ratio of Eevee-spCDK and the fluorescence intensity of Cdc13-mScarlet-I in individual cells (Figure 1F). This correlation was further confirmed by plotting the FRET/CFP ratio and the intensity of Cdc13-mScarlet-I as a function of cell length, a proxy for cell cycle progression in fission yeast (Figure 1G). In particular, a strong drop in FRET signal was detected in the largest cells, which correspond to the cells that have completed mitosis but have not undergone cytokinesis. To confirm that the FRET/CFP ratio reflects the phosphorylation of Eevee-spCDK, we substituted the ten possible phosphorylation serine or threonine sites with alanine residues, yielding Eevee-spCDK-10A (Figure 1H). As expected, cells expressing Eevee-spCDK-10A exhibited constant low FRET/CFP ratio values regardless of the intensity of Cdc13-mScarlet-I or cell length (Figures 1H and 1I). Note that Eevee-spCDK-10A was expressed under the stronger *tdh1* promoter^37^, because its expression level under the *adh1* promoter was insufficient for live-cell observation.

**Figure 1.**
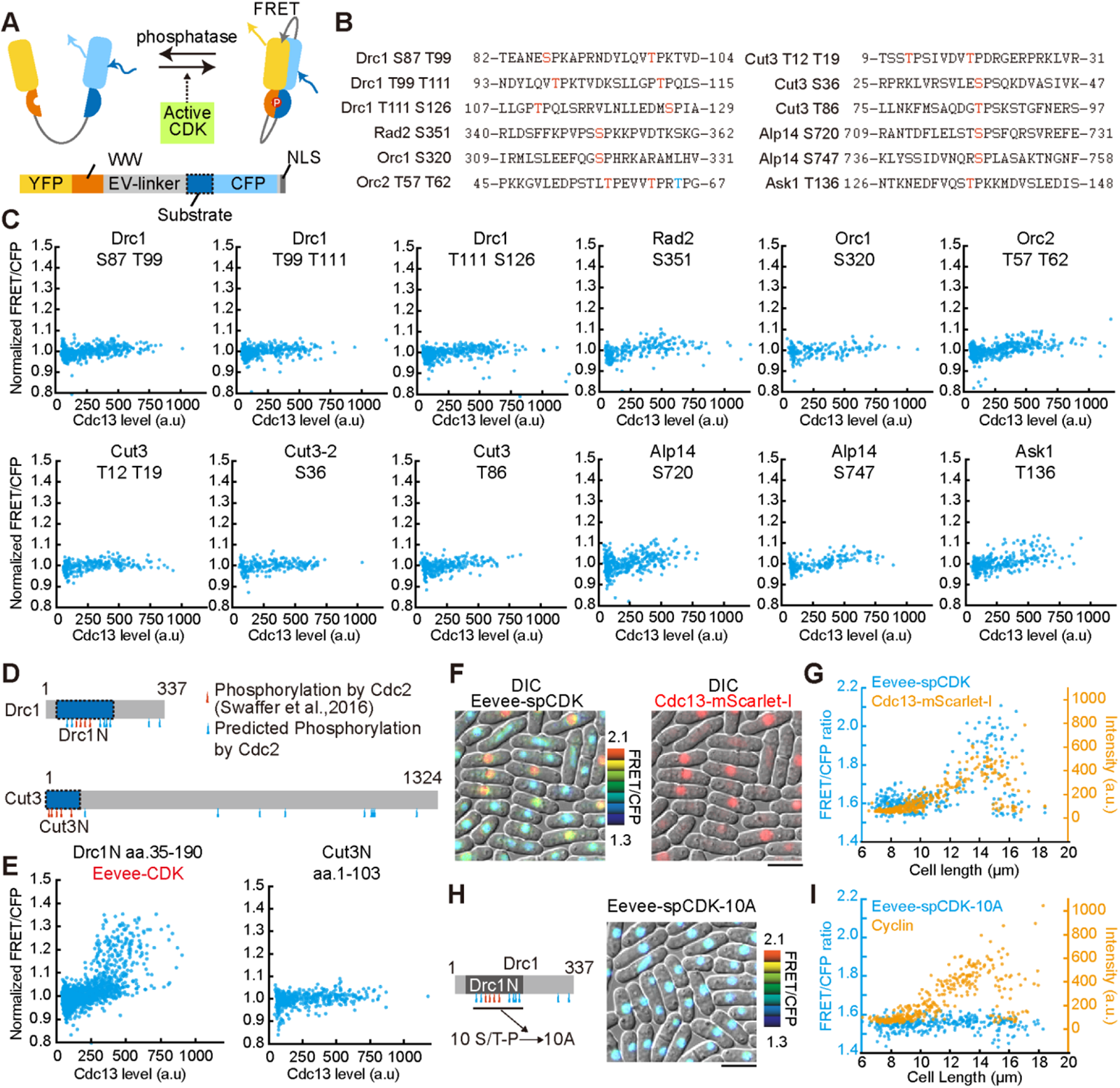
Development of a fission yeast CDK sensor, Eevee-spCDK. (A) Schematic illustration of the intramolecular FRET biosensor based on the Eevee backbone. Both cDNAs for fluorescent proteins (YFP: YPet; CFP: Turquoise2-GL) were optimized for fission yeast codons. NLS is fused to the C-terminus to monitor nuclear CDK activity. (B) Amino acid sequences of the 12 candidate peptides for the sensor domain of the CDK FRET biosensor. Red-colored residues indicate CDK-dependent phosphorylation sites reported in a previous phospho-proteomic analysis. Blue-colored residues indicate a potential CDK consensus sequence, S/T-P. (C) Quantification of the FRET/CFP ratio for the FRET biosensors containing the indicated sensor domains against cyclin (Cdc13-mScarlet-I) levels. Each dot represents a single cell. The FRET/CFP ratio was normalized by the median value in each experiment. (D) Schematic illustration of CDK phosphorylation sites in Drc1 and Cut3. Red arrowheads indicate CDK-dependent phosphorylation sites reported in previous phospho-proteomic analyses. Blue arrowheads are potential CDK consensus sequences, S/T-P. The dark blue region indicates the fragment used as sensor domain for the CDK FRET biosensor. (E) Scatter plots of the FRET/CFP ratio for the FRET biosensor containing the indicated sensor domains against cyclin (Cdc13-mScarlet-I) levels. Each dot represents a single cell. The FRET/CFP ratio was normalized by the median value in each experiment. (F) Representative images of fission yeast cells expressing Eevee-spCDK. The FRET/CFP ratio is visualized in the intensity-modulated display mode with pseudo color (left). The ratio image is overlaid with the corresponding DIC image. Endogenous cyclin levels (Cdc13-mScarlet-I) are shown in the red color channel and overlaid with the corresponding DIC image (right). Scale bar, 10 μm. (G) Scatter plot of the FRET/CFP ratio of Eevee-spCDK (cyan dots) and cyclin intensity (orange dots) against cell length. Each dot represents a single cell. (H) Schematic illustration of the non-phosphorylatable mutant of the Drc1 domain used in Eevee-spCDK-10A. The ten CDK target sites (S/T-P) are mutated to alanine (left). A representative image of fission yeast cells expressing Eevee-spCDK-10A is shown (right). Scale bar, 10 μm. (I) Scatter plot of the FRET/CFP ratio of Eevee-spCDK-10A (cyan dots) and cyclin intensity (orange dots) against cell length. Each dot represents a single cell.

### Characterization of Eevee-spCDK

Next, we evaluated whether Eevee-spCDK acts as a reliable CDK biosensor throughout the cell cycle. First, we confirmed that Eevee-spCDK expression under the control of the constitutive *adh1* promoter does not alter cell growth, while the excessive expression of either Eevee-spCDK or Eevee-spCDK-10A with the *tdh1* promoter, a stronger constitutive promoter than *adh1*, appeared to slightly reduce population growth (Figure S1A). Second, we validated the specificity and dynamic range of Eevee-spCDK by using an analog-sensitive mutant of Cdc2, *cdc2^as^*^11, 38^. Cells expressing *cdc2^as^* demonstrated a significant decrease in FRET/CFP ratio when treated with the non-hydrolysable ATP analog 1-NM-PP1 (Figure 2A), indicating that Eevee-spCDK indeed detects CDK activity. Furthermore, releasing these cells by washing out the 1-NM-PP1 inhibitor restored the FRET/CFP ratio with a typical time constant of approximately 20 min (Figure 2A). Next, *cdc2^as^* cells were arrested in late G2 using 3 h treatment with 1 μM 1-NM-PP1 and subsequently released from the G2 block into medium containing various concentrations of 1-NM-PP1 (Figure 2B). As a result, cells exhibited a wide range of FRET/CFP ratios that correlated with the concentration of the inhibitor in the release medium (Figure 2B). The fitted IC_50_ and Hill coefficient values were 9.2 ×10^2^ ± 7.8 × 10^1^ nM and 2.2 ± 4.0 × 10^-1^, respectively. Of note, we confirmed that the observed maximum FRET/CFP ratio was not saturated, as deletion of *dis2*, a catalytic subunit of PP1 (one of the main mitotic phosphatases in eukaryotes^39^), further increased the FRET/CFP ratio at the peak compared to WT cells (Figures S1B and S1C). Taken together, these results led us to conclude that Eevee-spCDK converts its phosphorylation status, which results from the combined activities of CDK and its counteracting phosphatase(s) (for simplicity, we will refer to this overall balance as “CDK activity”), into detectable FRET/CFP ratios in living fission yeast cells.

Taking advantage of the new FRET biosensor, we performed time-lapse experiments, following both CDK activity and endogenous cyclin levels (Cdc13-mScarlet-I) in living fission yeast cells. Consistent with the results of the snapshot population analysis (Figure 1G), we found that CDK activity gradually increases along with the accumulation of cyclin, and then decreases rapidly and concomitantly with chromosome segregation and cyclin degradation at mitotic exit (Figure 2C; Movie S1). Heatmap visualization of these experiments highlight the robust pattern of CDK activity dynamics, despite a small number of cells showing exceptionally short or long cell cycle durations (Figure 2D). The shapes of the averaged time courses for the FRET/CFP ratio and the expression level of Cdc13 (Figure 2E) were consistent with those derived from the snapshot images (Figure 1G). Interestingly, we recorded a small but detectable peak in CDK activity around 30 min after mitotic exit, when Cdc13 is expressed at its lowest level (Figure 2E). This small peak of CDK activity was also evident for small cells in the dispersed distribution of the FRET/CFP ratio values in Figure 1G. The timing of this peak coincided with S phase, as visualized by the aggregation of PCNA (Pcn1-mScarlet-I) (Figure 2F; Movie S2), suggesting that our biosensor is sensitive enough to detect the increase in CDK activity that is required for triggering S phase. Next, we compared the CDK activation dynamics obtained by Eevee-spCDK with those obtained by a previously reported translocation-based CDK activity reporter, synCut3^32^ (Figure 2G). SynCut3 translocates from the cytoplasm to the nucleus upon phosphorylation by CDK: the ratio of synCut3 fluorescence intensity between these two compartments (Nucleus/Cytoplasm; N/C) can be used to evaluate CDK activity. We found that the N/C ratio of synCut3 increases drastically at the G2/M transition but remains at a basal level in other cell cycle stages (Figure 2G). In contrast, Eevee-spCDK exhibits a more gradual increase in CDK activity through the cell cycle, with a rapid burst in activity around the time at which cells enter mitosis (Figure 2G). These data indicate that Eevee-spCDK has a higher sensitivity to CDK activity than synCut3, allowing the detection of weak CDK activity levels in fission yeast cells, even during the G1, S and G2 phases of cell cycle.

Next, we used Eevee-spCDK to visualize stress-induced changes in CDK activity, as stress signals are thought to generally modulate CDK function in order to delay cell cycle progression. Cells exposed to a hypertonic solution containing 400 mM of potassium chloride exhibited a rapid decline in CDK activity, typically within 5 min (Figure 2H; Movie S3). Although cell cycle progression was paused, Cdc13-mScarlet-I kept accumulating (Figure 2I; Figure S2A). Despite the maintenance of the hypertonic stress, CDK activity was eventually restored, approximately 30 min after adding KCl, and cell cycle progression resumed (Figure 2H). As anticipated, Eevee-spCDK-10A did not show any change in the FRET/CFP ratio during the osmotic stress (Figure 2H; Figure S2A). A decrease in CDK activity was also observed in cells transiently exposed to 1 mM hydrogen peroxide (Figures 2J and 2K; Figure S2B; Movie S4). In this case, we adopted a pulsed treatment, as most cells cannot resume vegetative growth under sustained oxidative stress. In this context, however, CDK activity did not recover as rapidly (Figure 2J), although Cdc13-mScarlet-I continued to accumulate (Figure 2K; Figure S2B). Indeed, we recorded an increase in CDK activity and a return to growth only 120 min after the transient oxidative stress (Figures 2J and 2K; Figure S2B). As with KCl treatment, no significant changes were observed with the Eevee-spCDK-10A system (Figure 2J; Figure S2B). We conclude that Eevee-spCDK is a useful and specific tool to monitor CDK activity at the single-cell level when fission yeast cells are exposed to various conditions.

**Figure 2.**
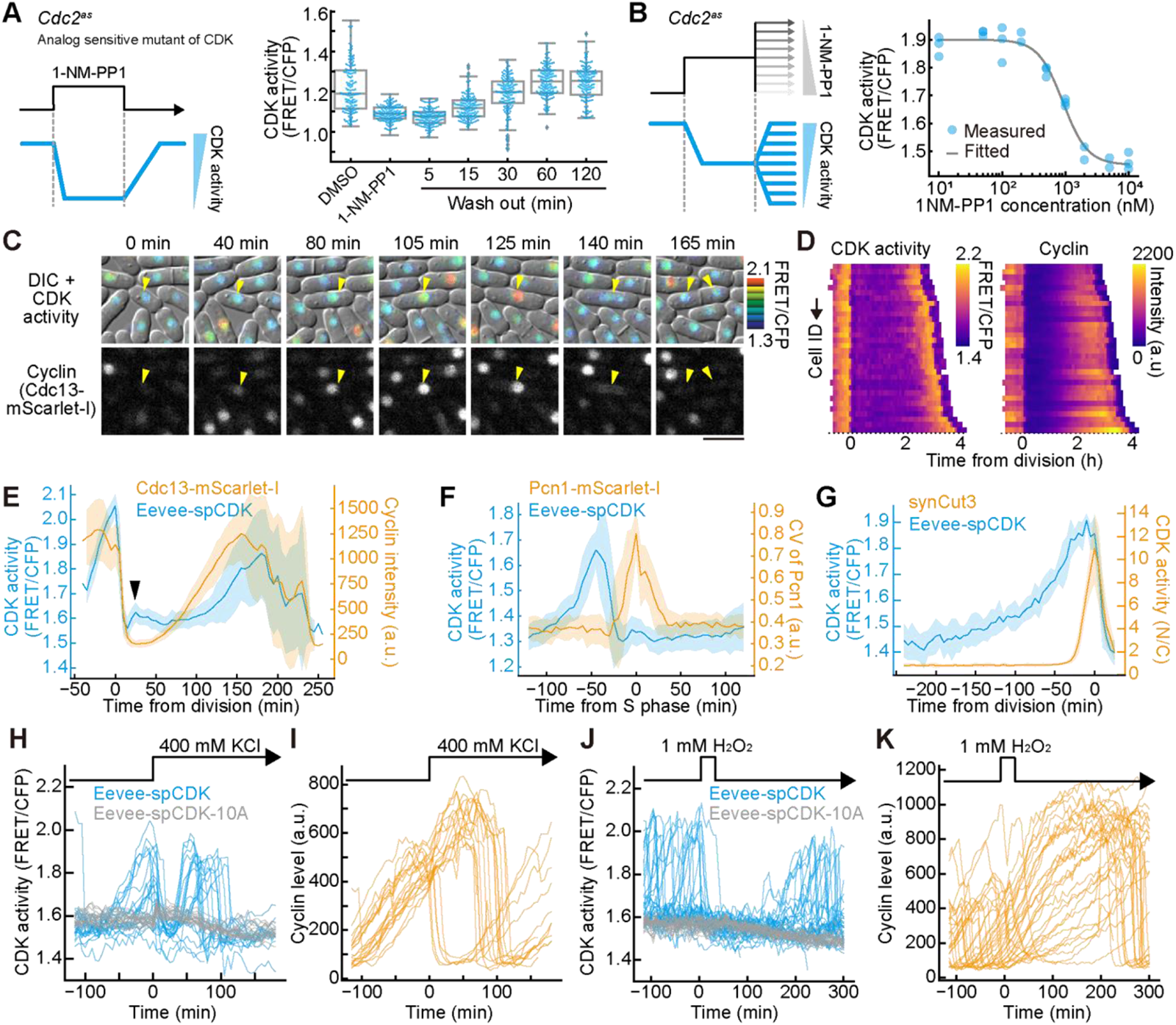
Characterization of Eevee-spCDK in the cell cycle of fission yeast. (A) Quantification of CDK activity in cells expressing an analog-sensitive mutant of Cdc2 (*cdc2^as^*) upon transient exposure to a non-hydrolysable ATP analog (1-NM-PP1). The left panel illustrates the schematics of the experimental procedure, and the right panel shows the quantification of the experimental results. Each dot represents the FRET/CFP ratio of an individual cell in the acquired snapshot images. The boxplots show the quartiles of the accumulated data with whiskers denoting the minimum and maximum except for the outliers detected as 1.5 times the interquartile range. n = 150 cells for each condition. (B) Dose–dependent response of G2-arrested cells expressing *cdc2^as^* when released in media containing various concentrations of 1-NM-PP1. The left panel illustrates the schematics of the experimental procedure, and the right panel shows the quantification of the experimental results. Each blue dot represents the average FRET/CFP ratio from more than 25 cells in the snapshot images. The solid black line represents the fitted curve based on the Hill function (IC_50_, 9.2 x10^2^ ± 7.8 x 10^1^ nM; Hill coefficient, 2.2 ± 4.0 x 10^-1^). (C) CDK activity and endogenous cyclin levels in single cells. The FRET/CFP ratio images are overlaid with the corresponding DIC images. The bottom panels show Cdc13-mScarlet-I. Yellow arrowheads identify the nucleus of a single cell throughout the time-lapse. Time 0 represents the start of the observation. Scale bar, 10 μm. (D) Heatmap representation of time-courses for CDK activity (left, FRET/CFP ratio) and cyclin levels (right, Cdc13-mScarlet-I intensity). Each row represents a single cell trace across time. Cells were aligned with the FRET/CFP peak as time 0. (E) Time-courses for CDK activity and cyclin levels. Time 0 is defined as the time point of nuclear division. Lines show the mean CDK activity (blue) and cyclin level (orange) for 39 individual cells. Shaded areas indicate standard deviations. The black arrowhead indicates a slight increase in CDK activity. (F) Time-courses for CDK activity and heterogeneity of Pcn1 distribution in the nucleus. The heterogeneity of Pcn1 in the nucleus was quantified. The coefficient of variation (CV) value of nuclear Pcn1 signal increases during S phase, as Pcn1 forms nuclear foci during DNA replication. Time 0 is defined as the time point when the CV value for Pcn1 reaches its maximum. Lines show the mean CDK activity (blue) and CV of Pcn1-mScalet-I (orange) from 13 cells, respectively. Shaded areas indicate standard deviations. (G) Time-course of CDK activity measured by Eevee-spCDK and synCut3. Time 0 is defined as the time point when the ratio of nuclear to cytoplasmic (N/C) synCut3 signals reaches its maximum. Lines show the mean FRET/CFP ratio of Eevee-spCDK (blue) and N/C ratio of synCut3-mScarlet-I (orange) for 9 individual cells. Shaded areas indicate standard deviations. (H and I) Response of CDK activity (H) and cyclin levels (I) to osmotic shock (400 mM KCl). Time 0 is defined as the time point when KCl was added. The data are shown in a line plot for 20 cells showing high cyclin levels during the 30 min prior to the stress (all data are shown in Figure S2A). (J and K) Response of CDK activity (J) and cyclin levels (K) to oxidative stress (1 mM H_2_O_2_ for 30 min). Time 0 is defined as the time point when H_2_O_2_ was added. The data are shown in a line plot (n = 31 cells; all data are shown in Figure S2B).

### Characterization of CDK activity dynamics and cell cycle events at the G2/M transition in fission yeast

During the analysis of the time series for evaluating CDK activity dynamics, we reproducibly observed a biphasic behavior: CDK activity increases gradually in the earlier part of G2 and more abruptly in a later phase (Figure 3A, point #7). We will refer to the point at which this rate change occurs as the “change-point”. In previous theoretical research, this switch was proposed to be the result of the Wee1/Cdc25-mediated feedback loops and to correspond to the G2/M transition^40, 41^. However, unlike with mammalian cells, it remains technically difficult to precisely define the G2/M transition of fission yeast because of the absence of nuclear envelope breakdown. Therefore, we estimated the timing of mitotic onset by monitoring key cell cycle events that occur around the G2/M transition. Specifically, we used the following markers (Figure 3A and 3B): cyclin degradation (anaphase onset, point #1, – 6.2 min from nuclear division), CDK activity drop (mitotic exit, blue point #2, – 8.1 min), cell growth halt (M phase, point #3, – 10.3 min), peak of CDK activity (M phase, point #4, – 17.8 min), spindle emergence (early M phase, #5, – 18.5 min), peak of cyclin level (point #6, – 37.0 min), and change-point (point #7, – 54.6 min). Of note, the peak of cyclin level precedes cyclin degradation by 30.8 min because cyclin localizes to the spindle pole body (SPB) before M phase entry^42^, translocating cyclin to a different imaging focal plane and causing an apparent decrease in averaged fluorescence brightness in the nucleus. Since spindle emergence is an early M phase event, we conclude that the G2/M transition occurs between the cyclin peak and spindle appearance (Figure 3B). In addition, the change-point of CDK activity precedes the cyclin peak by 17.6 min, suggesting that the change in CDK activity dynamics may be critical in late G2, prior to the G2/M transition.

**Figure 3.**
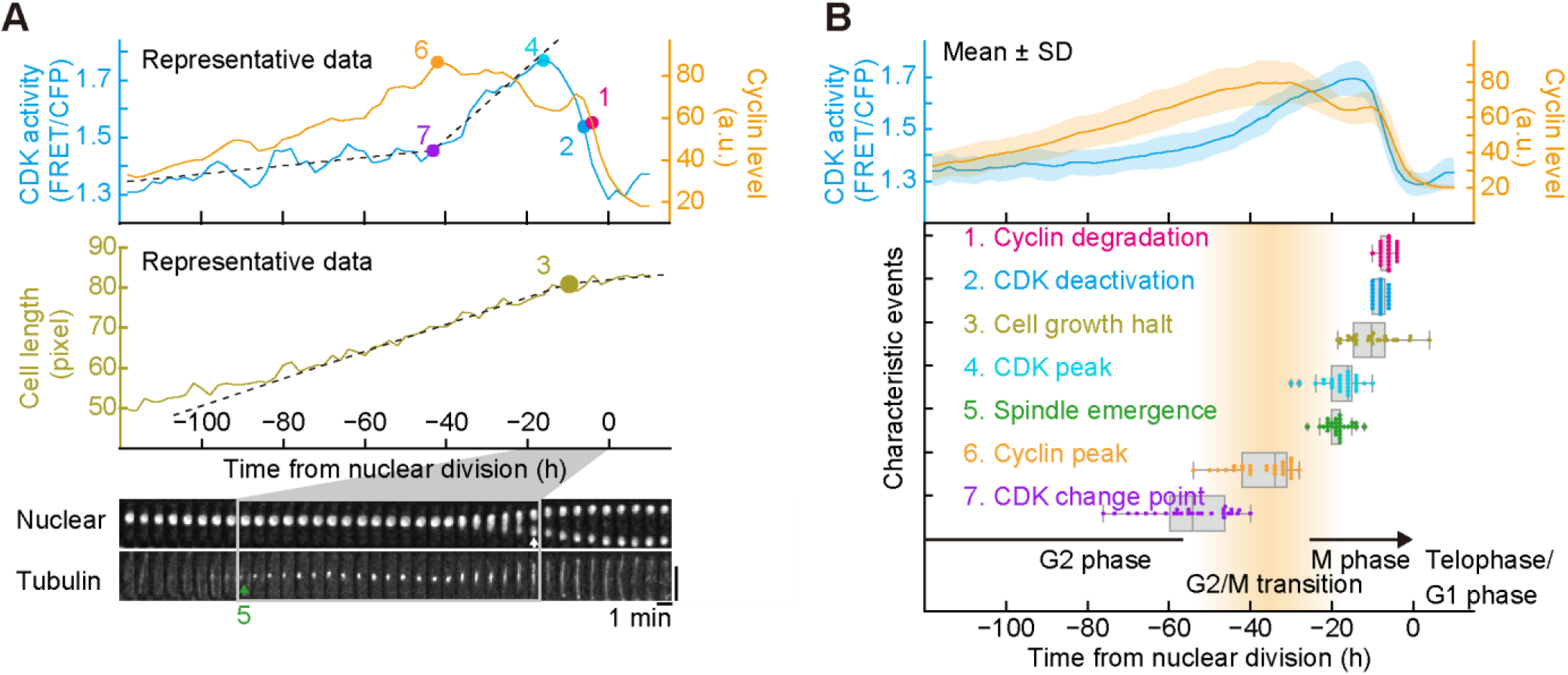
Timing of characteristic events occurring around the G2/M transition. (A) Definition of the cell cycle events that we monitored around the G2/M transition. Representative time courses for CDK activity and cyclin level (top) as well as cell length (middle) are shown. The bottom panel displays a montage of images showing the nucleus (Eevee-spCDK) and tubulin (mCherry2-atb2 under the constitutive *adh15* promoter) for a single cell throughout the G2/M transition. #1, cyclin degradation (the steepest negative slope in cyclin level); #2, drop in CDK activity (the steepest negative slope in CDK activity); #3, cell growth halt (cross point of the fitted two linear functions for cell length); #4, peak in CDK activity (the time point when CDK activity reaches its maximum); #5, spindle emergence (first visible spindle in time-lapse images); #6, peak in cyclin level (the time point when cyclin level reaches its maximum); #7, change-point (cross point of the fitted two linear functions for CDK activity). Time 0 is defined as the time point of nuclear division. Vertical scale bar, 10 μm. (B) Averaged time courses (top) for CDK activity (blue) and cyclin level (orange), and quantification of the timings of the events defined in A (bottom). Each single dot represents the data of an individual cell. The boxplot shows the quartiles of the accumulated data with whiskers denoting the minimum and maximum except for the outliers detected as 1.5 times the interquartile range. n = 27 cells for #1, #2, #3, #4, #6, and #7. n = 29 cells for #5.

### Decoupling CDK activity from cyclin accumulation in *pom1Δ* daughter cells

Live-cell imaging with Eevee-spCDK revealed similar dynamics between CDK activity and cyclin expression (Figures 2E and 3). To further investigate the relation between these two parameters during cell cycle progression, we used fission yeast cells lacking the *pom1* gene (*pom1Δ*). Pom1 is a mitotic inhibitor, the gradient of which was proposed to account in part for the coupling of cell size and cell cycle progression in *S. pombe*^17, 18^, although the original model has been more recently challenged^19–21, 43, 44^. Pom1 is also involved in determining the position of septation between the two future daughter cells by preventing contractile ring formation at the cell tip^45^. Consistent with previous reports^46^, the division plane in *pom1Δ* cells appeared to be asymmetric (Figure 4A). We defined an asymmetry index and found that *pom1Δ* cells showed a higher value than WT cells (Figure 4B). Overall, the peak of CDK activity in *pom1Δ* cells was quantitatively almost identical to that in WT cells (Figure S3A).

Using *pom1Δ* cells, we compared pairs of daughter cells that showed different lengths at birth as a result of asymmetric division. Thus, cells with an asymmetry index exceeding 0.25 were manually selected (Figure 4B, dashed line) for further investigation. Cells displaying a short size at birth showed a prolonged cell cycle duration compared to the long ones (Figure 4C). Nevertheless, short cells eventually entered mitosis with a shorter length at division (determined at the time of septum formation) than long cells (Figure 4D). We note that the net growth of cell length within one cell cycle demonstrated a slight negative correlation between short and long cells, consistent with a previous report^44^ (Figure S3B). However, this was not sufficient to compensate for the asymmetry in cell length at birth (Figure 4D; Figure S3B). These data suggest a less efficient size homeostasis by deleting *pom1*, consistent with a recent study^47^.

Next, we analyzed cyclin levels and CDK activity during the cell cycle of short and long cells in the *pom1Δ* background. The scatter plot of cyclin levels and CDK activity that we obtained indicated that long cells display the same maximum CDK activity than small cells but with lower amount of cyclin (Figure 4E). Of note, both short and long cells entered S phase immediately after mitotic exit (Figure S3C), suggesting that the G2 phase accounts for the uncouple of CDK activity from cyclin level. In addition, the short and long cells accumulated cyclin at similar rates (Figure 4F, left), but the short cells took a longer time to complete the cell cycle than the long cells (Figure 4C), resulting in higher amounts of cyclin at the cyclin peak just prior to the G2/M phase transition (Figure 4F, right). In contrast, CDK activity in short vs. long cells showed similar patterns, *i.e.* low basal activity just after mitosis and steep activation before mitotic onset (Figure 4G, left), although cell cycle duration was different between the two types of cells (Figure 4C). Consistent with this and with our conclusions from the scatter plot, there was no significant difference in CDK activity at the peak of cyclin level just prior to the G2/M phase transition between short and long cells (Figure 4G, right). This suggests that post-translational modification of CDK play a critical role in this situation. In conclusion, our approach demonstrated that CDK activity is not always coupled to cyclin levels, indicating that depending on the context, the relative importance of the different inputs on CDK function may vary. Eevee-spCDK is therefore a useful tool for visualizing the contribution of the different branches of the cell cycle control network to cell proliferation, including cyclin binding and post-translational modification of CDK.

**Figure 4.**
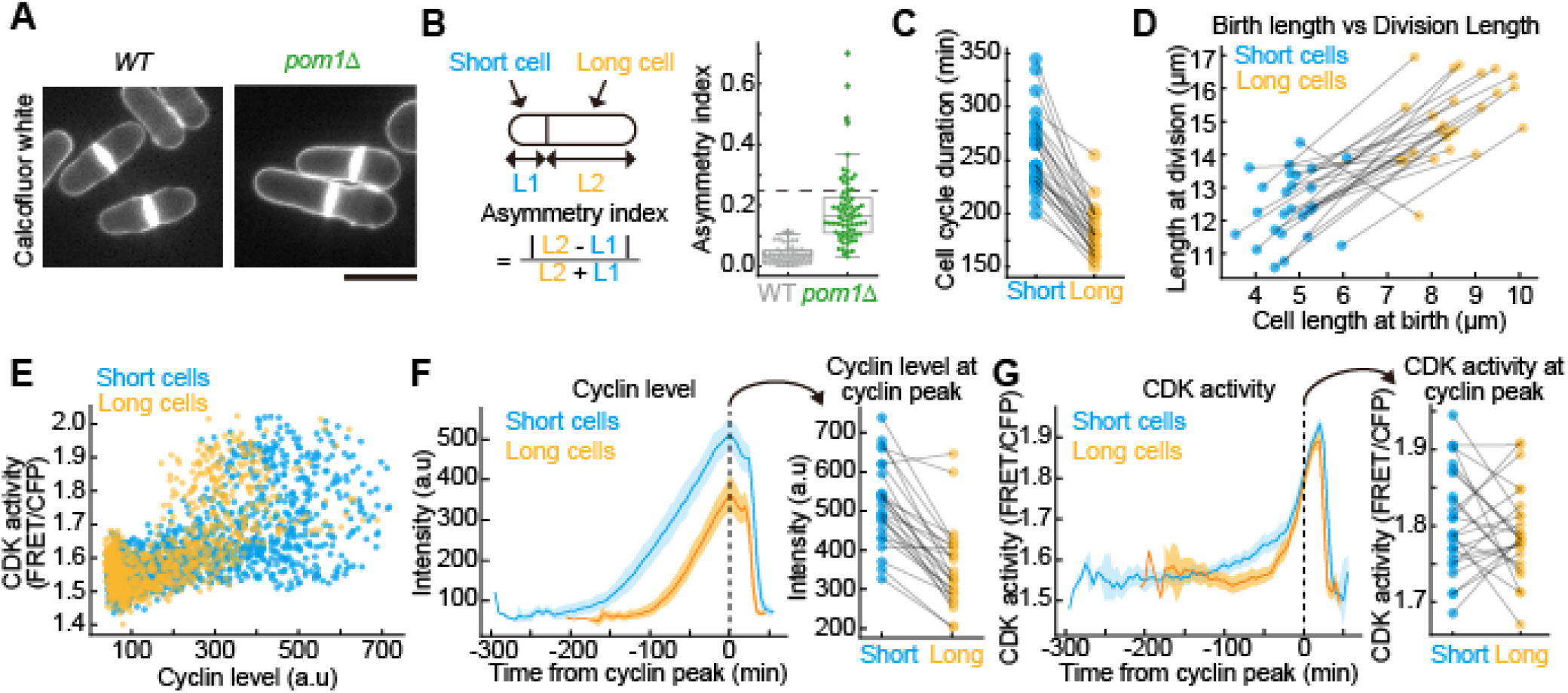
Decoupling CDK activity from cyclin accumulation in *pom1Δ* daughter cells. (A) Representative image of wildtype cells (WT) and Pom1-depleted cells (*pom1Δ*) stained by Calcofluor. Scale bar, 10 μm. (B) Quantification of the asymmetry index in WT and *pom1Δ*. The left panel represents the schematic definition of the asymmetry index, and the right panel shows the quantified distribution. The boxplot shows the quartiles of the accumulated data with whiskers denoting the minimum and maximum except for the outliers detected as 1.5 times the interquartile range. n = 74 cells for WT, n = 73 cells for *pom1Δ*. Cells with an asymmetry index exceeding 0.25 (black dashed line) were used for the following analyses. (C) Cell cycle duration of *pom1Δ* cells classified by cell length at birth. Black lines connect sister cells. Cell cycle duration was determined as the time from one nuclear division to the next. n = 25 cells. (D) Relation between cell length at birth and cell length at division, when a septum is newly formed, plotted for short and long cells. Black lines connect sister cells. n = 25 cells. (E) CDK activity plotted against cyclin levels in short and long cells. Each dot represents a single point in the individual time series data from 25 cells. (F and G) Cyclin accumulation (F) and CDK activity dynamics (G) during the cell cycle in short and long cells. The left panel shows the averaged time series for 25 short (blue) and long (orange) cells, with a 95% confidence interval (shaded areas). Time 0 is defined as the time point of the cyclin peak. The right panel shows the level of the cyclin peak (F) and the CDK activity peak (G). Black lines connect corresponding sister cells.

### Monitoring the CDK activity threshold at the G2/M transition

In fission yeast, it has been proposed that the G2/M transition is triggered when CDK activity exceeds a certain threshold^13, 48^. Interestingly, it was later-on demonstrated using an analog-sensitive version of Cdc2 that this also applies to S phase onset, although at a much lower threshold than for mitosis^11^. Consistent with this idea but with a more direct proxy for CDK activity, we found that at the time cells commit to mitosis, short and long cells in the *pom1Δ* strain exhibit significant differences in cyclin levels (Figure 4F), but similar CDK activity (Figure 4G). To further evaluate the model of CDK activity thresholds during the cell cycle, we quantified CDK activity in cells overexpressing either *cdc25* or *wee1*, which are positive and negative regulators of CDK, respectively. To this end, we constructed a system enabling gene expression at various levels in fission yeast. This system, which is based on the plasmid for Stable Knock-In (pSKI)^49^, contains a series of promoters driving a wide range of expression levels (designated as pSKI-v2; Figure 5A). The current pSKI-v2 lineup includes the following promoters: the constitutive promoters *adh1*, *tdh1*, *act1*, *pak1*, *pom1*, and *rga3* (abbreviated as A1, T1, AC1, PA1, PO1, and R3, respectively), and the inducible promoter *nmt1* (abbreviated as N1). In addition, we reduced the strength of the A1, T1, and N1 promoters by mutating their TATA box. Gene expression levels were then assessed by quantifying the fluorescence intensity of mNeonGreen under the control of each promoter (Figure 5B; Figure S4A). The properties of some of the attenuated variants were assessed by Western blot (Figure 5C; Figure S4B). Thus, pSKI-v2 achieves a high and continuous dynamic range of expression levels of approximately 1,000-fold.

Next, we established 64 fission yeast strains expressing different levels of Cdc25 and/or Wee1 (Figure 5D) using the pSKI-v2. The expression levels of each enzyme were normalized by the expression level of endogenous Cdc25 or Wee1 (Figures S5A-S5E). We then mapped the average cell length at division (Figure 5E) and its coefficient of variation (CV) (Figure S5F) on a two-dimensional plane of Cdc25 and Wee1 expression levels. Consistent with previous reports^50, 51^, the overexpression of Cdc25 and Wee1 led to a decrease (blue dots) and an increase (red dots) in cell size at division, respectively (Figure 5E). Excess overexpression of either Cdc25 or Wee1 resulted in cell death (indicated as cross marks in Figure 5E). As anticipated based on the fact that Wee1 and Cdc25 have opposite competing functions in controlling CDK phosphorylation on tyrosine 15, the permissive expression level of one gene increased with the degree of overexpression of the other, with the observed lethality resulting from a significant disequilibrium between Wee1 and Cdc25 levels (Figure 5E). Note that certain combinations of Wee1 and Cdc25 expression resulted in cells dividing at an average size similar to that of WT (Figure 5E), but with higher cell-to-cell heterogeneity (Figure S5F).

Using these strains, we next examined whether the overexpression of either Cdc25 or Wee1 perturbs the CDK activation pattern during the cell cycle. For simplicity, we only analyzed CDK activity in cells that overexpressed one of these factors (Figure 5F). As observed in short and long cells in the *pom1Δ* strain, CDK activity in Cdc25- or Wee1-overexpressing cells showed similar temporal patterns to wild type, including the rapid rise just prior to the onset of mitosis (Figure 5F; Figure S5G), although cell length at division and cell cycle duration were different between these strains (Figures 5E and 5F; Figure S5H). Next, we quantified the ratio of variance between the strains to variance within the strains, or *F*-value, as a proxy for the differences in CDK activity between Cdc25-overexpressing, Wee1-overexpressing, and WT cells. Strikingly, we found that the difference became less significant around the G2/M phase transition (Figure S5I), suggesting that mitotic entry in all three genetic backgrounds rely on comparable levels of CDK activity. This strongly supports the model of CDK activity threshold at mitotic onset. To further compare CDK activity levels in these distinct strains that exhibit different cell cycle durations (Figure 5F, top panels), we selected three time points as defined in Figure 2: the change-point (#7), around G2/M transition (between #6 and #5), and the peak of CDK activity (#4). Again, our analyses showed that CDK activity around the G2/M phase transition (F-value, 38.7) was less different among the three strains than at the change-point (F-value, 136.4) and peak in CDK activity (F-value, 69.7) (Figure 5G). Of note, this trend was also confirmed when comparing short and long *pom1Δ* cells (Figure S5J). Altogether, these results support the idea that there is an absolute threshold in CDK activity that defines the G2/M transition.

Finally, we further dissected CDK activity at the change-point, which precedes the G2/M phase transition by about 20 min. Based on previous mathematical studies^40, 41^, the change-point may correspond to a saddle-node bifurcation point, which triggers the positive feedback loops mediated by Cdc25 and Wee1. This bifurcation point leads to the hyperactivation of CDK at the G2/M transition. We also reproduced the CDK activation dynamics through mathematical modeling and numerical simulation (Figure S5K), revealing a change-point just prior to CDK hyperactivation^40, 41^. Together with the experimental results (Figure 5G, middle), we propose that Wee1 and Cdc25 modulate the level of CDK activity that is required for switching to the faster rate of CDK activity increase prior to mitotic onset. More specifically, our data imply that alteration of Wee1 and/or Cdc25 levels changes the CDK activity that is necessary to activate the double positive feedback loop on CDK function mediated by these two enzymes. For instance, overexpression of the CDK inhibitor Wee1 generates a negative imbalance in the system that can only be overturned by higher CDK activity than in WT cells. To investigate this idea, we plotted CDK activity before and after the G2/M phase transition upon changes in Wee1 and Cdc25 expression in the mathematical model^52^. The numerical simulation revealed a sharp increase in CDK activity when the CDK-cyclin complex reached a particular level, which seemed to correspond to the change-point (Figure 5H, purple dots). Consistent with the experimental data (Figure 5G, left), overexpression of Wee1 elevated the level of CDK activity required for the change-point (Figure 5H, orange lines; Figure 5I, upper panel). Conversely, numerical simulations using previously reported parameters^52^ indicated that overexpression of Cdc25 decreased the level of CDK activity required for the change-point (Figure 5H, green lines; Figure 5I, lower panel). However, our experimental results showed comparable CDK activity at the change-point between WT and Cdc25-overexpressing cells (Figure 5G, left). This discrepancy may be explained by the initial parameter values used for our simulation. Collectively, our results suggest that the relative strengths of the Wee1-mediated inhibition vs Cdc25-mediated activation of CDK tune the level of CDK activity that is required to activate the associated feedback loops, resulting in the hyperactivation of CDK at the onset of M phase. This also shapes the timing of the G2/M transition, as cyclin B accumulates at a constant rate over time (Figure 4F and Discussion).

**Figure 5.**
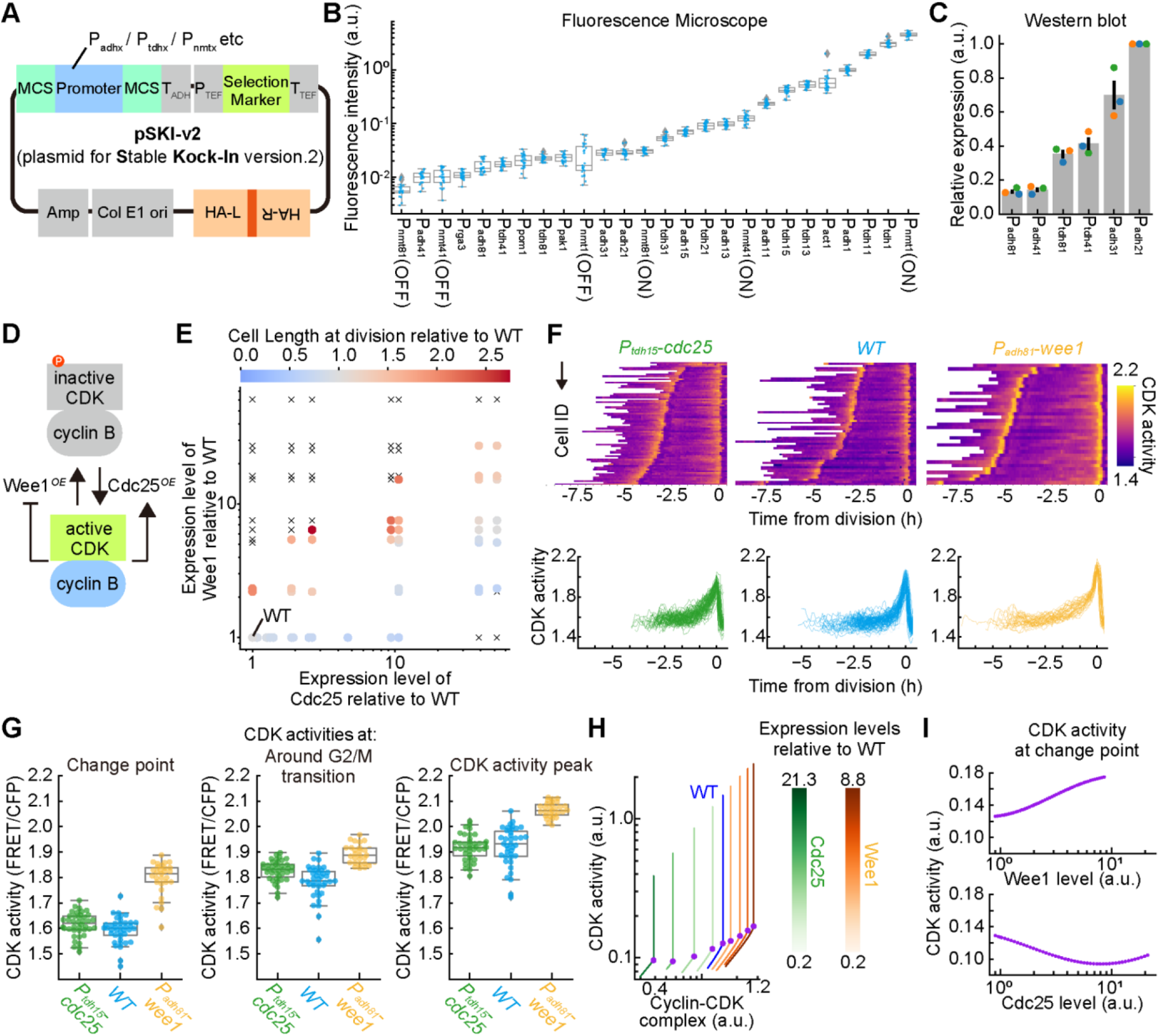
Effect of Cdc25 and/or Wee1 overexpression on cell length at division and CDK activity during the cell cycle. (A) Schematic illustration of the Stable Knock-In version 2 (pSKI-v2) plasmid. (B and C) Quantification of the promoter strength of pSKI-v2 plasmids. (B) Fluorescence intensity of mNeonGreen expressed under the indicated promoters was measured by fluorescence microscopy (Figure S4A). Each dot represents the signal intensity from a single cell. The boxplot shows the quartiles of data with the whiskers denoting the minimum and maximum except for the outliers detected as 1.5 times the interquartile range (n > 20 cells for each strain). (C) The protein amounts of mNeonGreen-3xFLAG were measured by Western blotting (Figure S4B) for the indicated weak promoters, for which fluorescence measurements were not reliable. Each dot represents a quantification from triplicate experiments. Error bars denote the standard error. (D) Schematic representation of the experiments modulating CDK activity control through the inhibitory phosphorylation of CDK by overexpressing *cdc25* and/or *wee1*. The activating phosphatase Cdc25 and the inhibitory kinase Wee1 regulate the inhibitory phosphorylation of CDK. (E) Cell length at division in mutant strains expressing *cdc25* and/or *wee1* at various levels. The levels of Cdc25 and/or Wee1 are normalized by that in WT cells (Figure S5A-5E). Each dot or cross mark corresponds to a strain harboring a specific combination of promoters for overexpressing *wee1* and *cdc25* at different levels. Cross marks represent combinations that lead to lethality. The relative cell length to WT in each strain is color-coded as shown at the top. n > 20 cells for each strain. (F) Time series for CDK activity in Cdc25-overexpressing (green), WT (blue), and Wee1-overexpressing (orange) cells. Upper panels and lower panels represent heatmap and line plots, respectively. Each row in the heatmaps and single solid line in the line plots correspond to an individual cell. n = 81 for Cdc25-overexpressing, n = 82 for WT, and n = 33 for Wee1-overexpressing cells. (G) CDK activity at the change-point (left), around the G2/M phase transition (middle), and CDK activity peak (right). The boxplot shows the quartiles of the accumulated data with whiskers denoting the minimum and maximum except for the outliers detected as 1.5 times the interquartile range. Data comes from the time series in F. (H) Numerical simulation of CDK activity around the G2/M phase transition. The values of CDK activity for a range of cyclin-CDK levels were calculated. The blue line represents the results for WT. The green and orange lines represent the results for Cdc25- and Wee1-overexpressing cells, respectively. The color intensity of the lines show the degree of overexpression, as shown in the color bars on the right. Purple circles indicate the time point when CDK activity abruptly increases (change-point). (I) Relation between the CDK activity required for the change-point and the expression of either Cdc25 or Wee1 from the mathematical model, as described in H.

### Monitoring CDK activity in mammalian cells

The fact that human *CDK1* complements fission yeast *cdc2*^53^ prompted us to examine whether Eevee-spCDK also allows for monitoring CDK activity in mammalian cells. To this end, we established a HeLa cell line stably expressing Eevee-spCDK with mCherry-hGeminin as an S/G2/M phase marker^54^ and H2B-iRFP as a nuclear marker (Figure 6A; Figure S6A; Movie S5). The cells were then imaged using a wide-field fluorescence microscope to quantify the temporal trajectories of the FRET/CFP ratio of Eevee-spCDK and mCherry-hGeminin intensity during the cell cycle. Each trajectory was aligned with the end of mitosis as time 0, as determined by the decrease in mCherry-hGeminin intensity (Figure 6B; Figure S6B). The FRET/CFP ratio of Eevee-spCDK exhibited a sudden increase at the onset of M phase, followed by an abrupt drop at mitotic exit. These dynamics are reminiscent of the typical cyclin B-CDK1 activity pattern previously described in mammalian cells^26^. In addition to this transient increase in the FRET/CFP ratio at G2/M, we detected a preceding step-like increase 10 h before mitotic onset. When the time-trajectories of the FRET/CFP ratio of Eevee-spCDK and mCherry-hGeminin were aligned with the onset of Geminin accumulation as time 0, which corresponds to S phase onset, the step-like increase in FRET/CFP ratio started about 1 h before the G1/S phase transition (Figure 6C). Therefore, this change in activity might reflect the activation of Cyclin E/A-CDK2, which is known to be required for entry into S phase and DNA replication^55, 56^.

Finally, to examine the specificity of Eevee-spCDK, HeLa cells expressing Eevee-spCDK were treated with CDK subtype-specific inhibitors (Figure 6D). CDK1 and/or CDK2-directed inhibitors decreased the FRET/CFP ratio, while neither DMSO nor the CDK4/6-specific inhibitor Palbociclib affected the signal (Figure 6D). We further divided the cells into two groups, a Geminin-high *vs* Geminin-low, and found that the Geminin-high group exhibited higher sensitivity to CDK1 and/or CDK2 inhibitors than the Geminin-low group (Figure S6C). These data show that CDK1/2 inhibitors reduce the FRET/CFP ratio only in cells that are in S, G2, or M, during CDK2 and CDK1 are known to be activated. Collectively, these results led us to conclude that Eevee-spCDK is compatible with the monitoring of CDK1/2 activity in mammalian cells.

**Figure 6.**
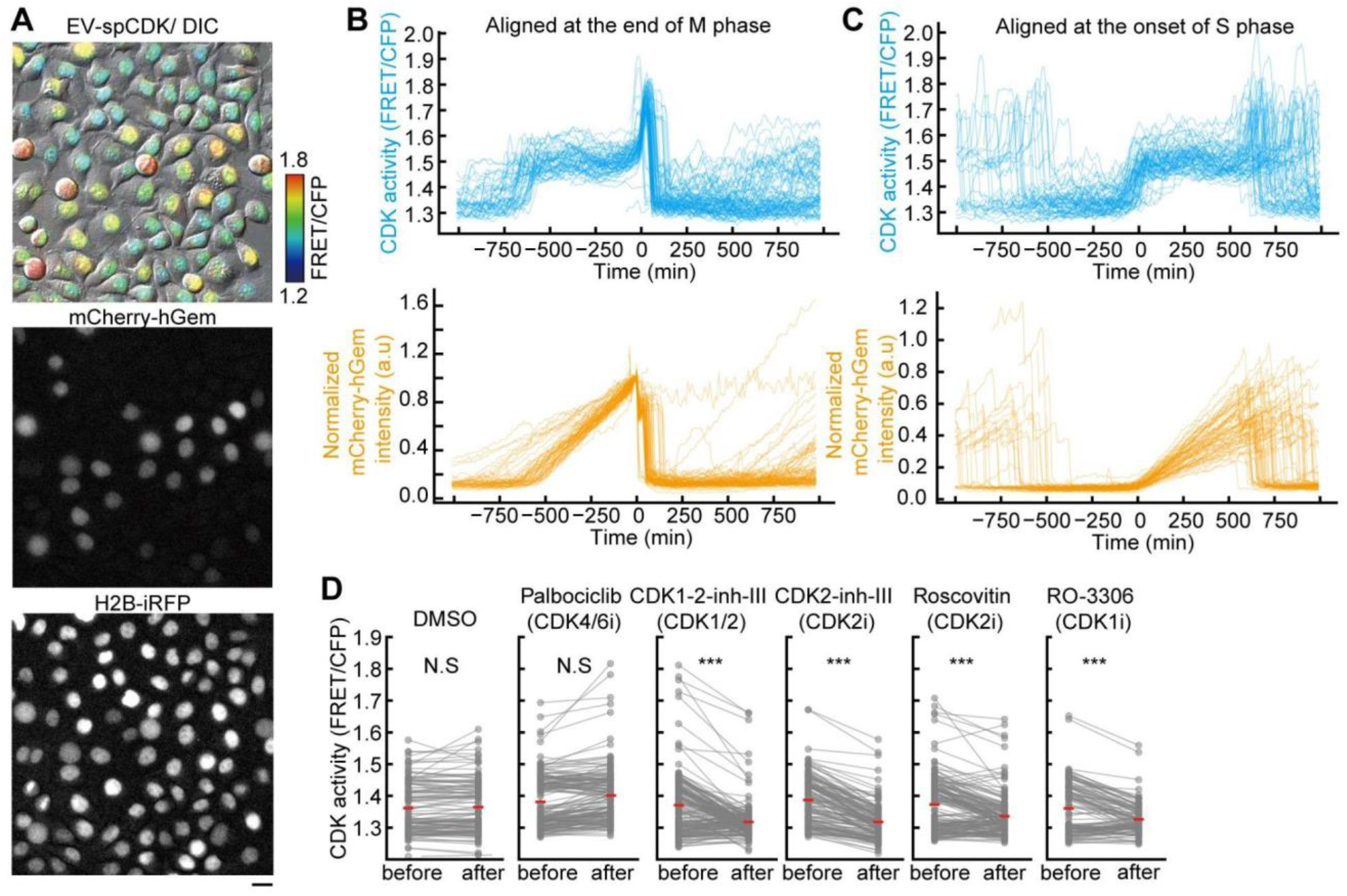
Monitoring CDK1/2 activity with Eevee-spCDK in mammalian cells. (A) Representative images of HeLa cells expressing Eevee-spCDK. FRET/CFP ratios are visualized in the intensity-modulated display mode with pseudo colors. The ratio image is overlaid with the corresponding DIC image (top). The signals for mCherry-hGeminin (middle) and H2B-iRFP (bottom) are shown in gray scale. Scale bar, 20 μm. (B and C) Quantification of CDK activity (FRET/CFP ratio) during the cell cycle. The time-series for CDK activity were aligned at the peak of mCherry-hGeminin intensity, indicating mitotic exit (B), or at the start of the increase in mCherry-hGeminin intensity, corresponding to the time point of S phase entry (C). Each line represents the trace of FRET/CFP ratio (Eevee-spCDK, blue) and mCherry-hGeminin intensity (orange) in a single cell. n = 85 cells and n = 70 cells, respectively. (D) Comparison of CDK activity (FRET/CFP ratio) before and after treatment with the indicated CDK inhibitors. Cells were treated with the inhibitors (10 μM) for 30 min. Each dot represents the FRET/CFP ratio from a single cell. The lines link the same cells before and after treatment. One-sided paired *t*-tests were performed. Not significant (N.S.), *p* > 0.05; ***, *p* < 0.001). n = 118, 157, 176, 140, 164, 115 cells for each panel.

## Discussion

In this study, we developed a novel CDK FRET biosensor, Eevee-spCDK, and monitored the dynamics of CDK activity—or the balance between CDK activity *per se* and related phosphatases—during the cell cycle in *S. pombe* and HeLa cells. In fission yeast, time-lapse imaging with Eevee-spCDK revealed that CDK activity first increases at the time of S phase. While it has been previously proposed^13^, our results provide the first experimental evidence for this transition in CDK activity levels as cells progress into DNA replication (Figure 2E). CDK activity then gradually increases during G2 and rapidly rises around the onset of mitosis. It remains high during M phase, and declines abruptly to basal levels following mitotic exit (Figure 7, blue line). The small increase in CDK activity during S phase resembles the step-like increase in CDK activity prior to S phase entry in HeLa cells (Figure 6). Given that little Cdc13 accumulates during G1 and S phase, other cyclins (Puc1, Cig1, Cig2) may be involved in this slight activity change.

**Figure 7.**
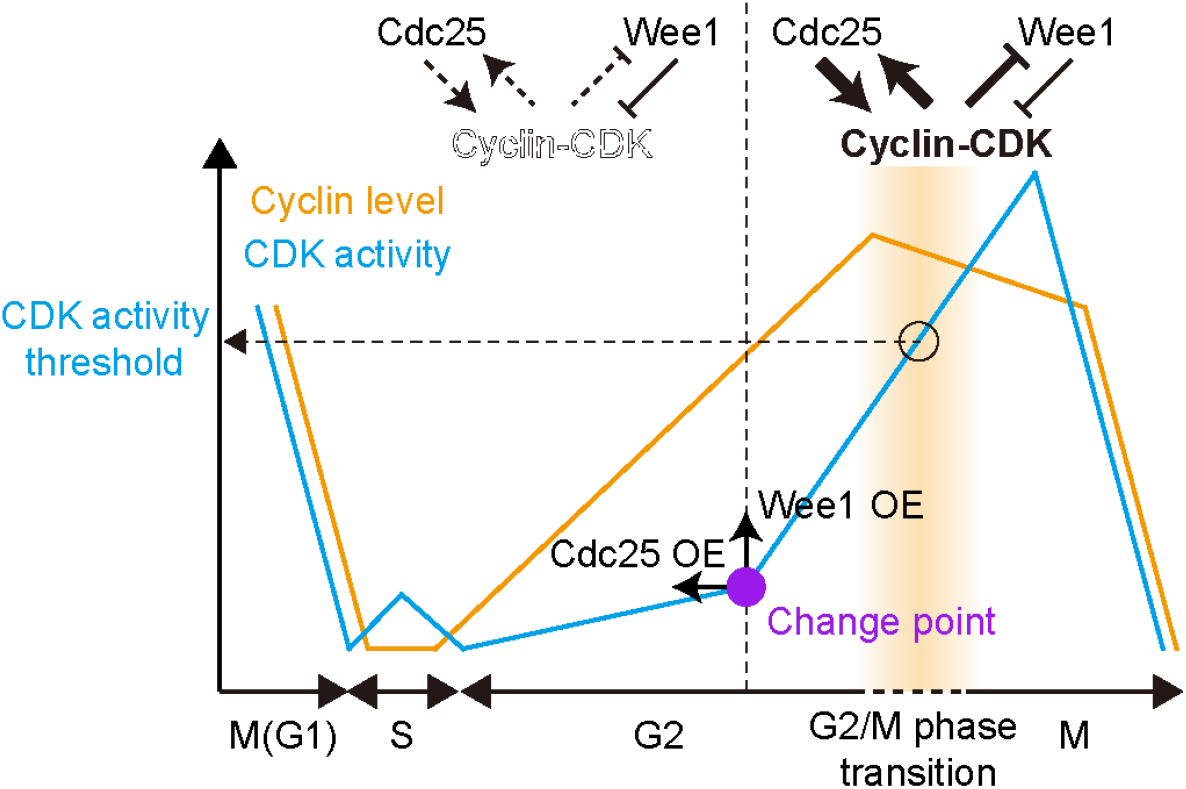
A model of the regulation of CDK at the G2/M transition. The typical dynamics of cyclin (orange) and CDK activity (blue) in fission yeast are shown.

Remarkably, our study also reveals that perturbations in cell length at birth (Figure 4) or in key regulators of CDK function (Figure 5) do not impact the levels of CDK activity that are associated with the G2/M transition (Figure 5G; Figure S5I). These observations provide strong and direct experimental evidence for the quantitative model of cell cycle control^11, 13, 48^. In a previous study, it was also suggested that a threshold in cell size-correlated cyclin B level may represent the potential molecular basis of cell-size homeostasis by linking division probability and cell size^16^. However, our results suggest that cyclin B level alone does not determine the threshold for the G2/M transition and hence the division timing. Even in cells of the same genetic background, cyclin B levels at mitotic onset can differ substantially, for instance between short and long *pom1Δ* cells (Figure 4F). Rather, cyclin B accumulates constantly and independently of cell length (Figure 4F) and cellular stress (Figures 2I and 2K). Consistent with these results, it has recently been reported that cyclin B acts as a timer-like molecule that monitors the time from cell division through accumulating at a constant rate. This may provide a basis for cell size homeostasis^47^. Cdc25 and Wee1 are also well-known key regulators of CDK impacting mitotic timing. Indeed, it has been demonstrated that the Cdc25/Wee1 protein ratio influences mitotic timing^22, 23^. Our present experiments corroborate this, showing that overexpression of Cdc25 or Wee1 changes the cell cycle length (Figure 5F; Figure S5H). In this context, it is noteworthy that Eevee-spCDK integrates the activities of CDK and phosphatases, as well as the effects of various regulations— such as post-translational modification of Cdc25 and Wee1— on CDK activity^15^.

The change-point, which delineates the biphasic increase in CDK activity during G2, is another important finding revealed by our live-cell imaging approach using Eevee-spCDK. Theoretical studies have predicted this biphasic pattern in CDK activity and the abrupt increase that we observe at the G2/M transition. This was proposed to result from the operation of the two positive feedback loops on CDK function mediated by Cdc25 and Wee1^40, 41, 52^. These positive feedback loops provide the system with bistability, adding hysteresis or a “point of no return” in cell cycle logic. What is the physiological meaning of the change-point? Our results using *pom1Δ* cells may provide a clue. In the *pom1Δ* background, the cells that were born short showed a higher CDK activity at the change-point and a longer cell duration, while the long ones exhibited lower CDK activity at the change-point and shorter cell cycle duration (Figure 4C; Figure S5J). This relation between CDK activity required for the change-point and cell cycle duration is also confirmed in Cdc25 or Wee1-overexpressing cells, which showed similar correlations between CDK activity at the change-point and cell cycle duration (Figures 5F and 5G; Figure S5H). These results led us to hypothesize that the CDK activity required for the change-point limits cell cycle duration. Because short and long cells share the same genetic background in our assay, the difference in CDK activity required for the change-point and cell cycle duration could be explained by the differences in cell size. Based on the fact that deletion of *pom1* may result in the loss of size-dependent regulation of Wee1, Cdc25 could be involved in the difference of CDK activity required for the change-point and cell cycle duration in *pom1Δ* cells. This model is supported by previous reports discussing the size-dependent expression of Cdc25^22, 23, 47^. Indeed, if short *pom1Δ* cells have less Cdc25 molecule, higher CDK activity would be required to overturn the bistability of the double positive feedback loops as it was suggested in the mathematical model (Figure 5H and 5I). Alternatively, the asymmetric distribution of Wee1 due to the size imbalance could explain the higher CDK activity at the change-point in long cells of *pom1Δ* strain, as shown in Wee1-overexpressing cells (Figure 5G). In any case, it would be needed to investigate a potential role for CDK activity required for the change-point in cell size homeostasis.

The advantage of Eevee-spCDK lies in its high sensitivity. While the previously reported CDK activity sensor in fission yeast, synCut3, detects the increase in CDK-mediated phosphorylation around M phase with a high S/N ratio^32^, Eevee-spCDK allows for monitoring CDK activity not only as cells enter mitosis, but also during S and G2 phases and perhaps even meiosis (which may only require the use of different promoter). In particular, the timing of the increase in N/C ratio of synCut3 coincides with the timing of the G2/M phase transition characterized by Eevee-spCDK (Figures 2G and 3B). Therefore, these two sensors can complement each other depending on the design of the experiment: synCut3 represents an easy and rapid tool to follow CDK activity at this particular phase of the cell cycle, whereas Eevee-spCDK enables more precise and quantitative monitoring of CDK activity throughout the cell cycle including S phase. However, the use of our biosensor requires relatively specialized equipment and dedicated analysis pipelines. The high sensitivity of Eevee-spCDK could be attributed to the nature of Drc1, known to act during S phase ^57^. In fact, the Drc1 fragment used for the sensor domain of Eevee-spCDK is phosphorylated from S phase^33^. On the other hand, one of the limitations of Eevee-spCDK is its specificity. Currently, it is unclear whether Eevee-spCDK distinguishes between the different qualitative CDK activities that rely on the binding to G1, S, and M phase cyclins in fission yeast. Furthermore, it should be noted that Eevee-spCDK has multiple phosphorylation sites, and the fragment we used harbors a total of 10 of these sites, 4 of which have been reported to be phosphorylated in fission yeast^33^. Experiments using analog-sensitive mutants of Cdc2 demonstrated that Eevee-spCDK quantitatively monitors CDK activity (Figure 2B). The multiple phosphorylation sites in Eevee-spCDK may thus confer the sensitivity and specificity of this biosensor to CDKs during the cell cycle, although the rationale for this remains elusive. In HeLa cells, the CDK2 sensor based on human DNA helicase B (DHB) showed a rapid increase at the timing of the G1/S phase transition, followed by a gradual rise during the G2 phase^29^. This gradual increase during G2 was not observed with Eevee-spCDK (Figure 6). This could be due to differences in the sensitivity and specificity for distinct qualitative cyclin/CDK pairs between the DHB-based CDK2 sensor and Eevee-spCDK.

In conclusion, our study provides a unique tool for monitoring nuclear CDK activity in live fission yeast cells. Interestingly, our CDK FRET biosensor, Eevee-spCDK, has the potential to allow for mapping the organization of CDK activity at the subcellular level by means of subcellular localization signals that can be used to tether the sensor to specific locations. Indeed, it has been reported that Cdc13 relocalizes from the nucleus to the SPB at the onset of M phase in fission yeast^42^. Similarly, cyclin B1 and B2 localize to different subcellular compartments in mammalian cells^58^. It remains poorly explored whether these changes are associated with more targeted modulation of CDK activity within cells. Monitoring CDK-dependent phosphorylation at the subcellular resolution may unravel extra layers of regulation that may be critical for cell cycle progression. Overall, Eevee-spCDK represents a breakthrough in deciphering the dynamic behavior of CDK activity in fission yeast based on the live imaging of single cells.

## Materials and Methods

### Plasmids construction

All plasmids constructed and used in this study are summarized in Table S1. The nucleotide sequences of mNeonGreen and Turquoise2-GL were optimized for codon usage in the fission yeast *S. pombe*^49^. For pSKI-v2, restriction enzyme sites (*Pvu*II, *Bam*HI, *Hind*III, *Spe*I) are inserted just before the promoter sequence to allow easy swapping of promoters. The promoters used had been previously studied^37^ and were subcloned by PCR amplification, followed by Gibson assembly with NEBuilder HiFi DNA assembly (New England Biolabs). Note that *P_tdh1_* used in this study is the shorter version of *Ptdh1*\* from a previous study^37^. As *P_adh1_* and *P_tdh1_* variants, the TATA-boxes in these two promoters (TATAAATA, -108 bp and -80 bp from the start codons of *adh1* and *tdh1*, respectively) were mutated to TATAAA for *P_adh11_*/*P_tdh11_*, TATAT for *P_adh13_*/*P_tdh13_*, TAAATATA for *P_adh15_*/*P_tdh15_*, TAAATA for *P_adh21_*/*P_tdh21_*, TAAAAATA for *P_adh31_*/*P_tdh31_*, AAAA for *P_adh41_*/*P_tdh41_*, and AT for *P_adh81_*/*P_tdh81_*^59^. For the screening of the FRET sensor, the candidate sequences shown in Figure 1B and Figure S1A-C were subcloned into a backbone plasmid using *Aor13*HI and *Not*I. To express Eevee-spCDK in mammalian cells, the cDNA of Eevee-spCDK was subcloned into pPBbsr-MCS, a PiggyBac transposon vector^35, 60^.

### ***S.*** *pombe* strains and cell culture

All *S. pombe* strains used in this study are summarized in Table S2. Unless otherwise stated, the standard methods and media selection for culturing *S. pombe* were based on those reported previously^61^. The transformation protocol was also a modified version of a previously reported protocol^62^. The *P_adh81_-wee1* cells were selected for the analysis in Figure 5F because *P_adh41_-wee1* cells were viable in the batch and plate cultures but ceased to grow after several rounds of divisions inside the microfluidic environment for unknown reasons.

### HeLa cell culture and stable cell line establishment

HeLa cells were a kind gift of Michiyuki Matsuda (Kyoto University). HeLa cells were cultured in Dulbecco’s Modified Eagle’s Medium (DMEM) high glucose (Nacalai Tesque) supplemented with 10% fetal bovine serum (FBS) (Sigma-Aldrich) at 37°C in a humidified atmosphere containing 5% CO2. For live-cell imaging, HeLa cells were plated on CELLview cell culture dishes (glass bottom, 35 mm diameter, 4 components: The Greiner Bio-One).

A lentiviral transgene system was applied to establish the HeLa cell line stably expressing mCherry-hGeminin and H2B-iRFP. For lentiviral production, HEK-293T cells were co-transfected with the pCSIIhyg-H2B-iRFP-P2A-mCherry-hGem, psPAX2, and pCSV-VSV-G-RSV-Rev by using Polyethylenimine ‘MAX’ MW 40000. Virus-containing media were collected 48 h after the transfection, filtered, and used to infect HeLa cells with 10 μg/ml polybrene. After the infection, cells were selected with 100 μg/mL hygromycin B (Wako, 085-06153) for at least one week, followed by single-cell cloning through a limited dilution method. To further introduce Eevee-spCDK, the PiggyBac transposon system was used. The pPBbsr-Eevee-spCDK was co-transfected with pCAGGS-hyPBase encoding the hyPBase PiggyBac transposase by using 293fectin^63, 64^. One day after the transfection, cells were selected with 20 μg/mL blasticidin S for at least one week. Subsequently, bulk cells with high expression of Eevee-spCDK were sorted and collected using an MA900 cell sorter (Sony).

### Microscopic imaging

Epifluorescence inverted microscopes (IX83; Olympus) were used for live-cell imaging. For spinning-disk confocal microscopy, the microscope was equipped with an sCMOS camera (ORCA Fusion BT; Hamamatsu Photonics), oil-immersion objective lenses (UPLXAPO 100X, NA = 1.45, WD = 0.13 mm and UPLXAPO 60X, NA = 1.42, WD = 0.15 mm; Olympus), and a spinning-disk confocal unit (CSU-W1; Yokogawa Electric Corporation). The excitation lasers and fluorescence filters used were as follows: excitation laser, 445 nm for Turquoise2-GL, 488 nm for mNeonGreen, and 561 nm for mScarlet-I; excitation dichroic mirror, DM405/488/561/640 for mNeonGreen, mScarlet-I, or YPet and DM445/514/640 for Turquoise2-GL; emission filters, 482/35 for Turquoise2-GL, 525/50 for mNeonGreen or YPet, and 617/73 for mScarlet-I (Yokogawa Electric). For wide-field fluorescence microscopy, the microscope is equipped with a Prime sCMOS camera (Photometrix) and oil-immersion lenses (UPLXAPO 100X and UPLXAPO 60X). The illumination settings and fluorescence filters were as follows: excitation wavelength, 390/15 for Calcofluor white, 438/24 for Turquoise2-GL, 475/28 for mNeonGreen, and 580/20 for mScarlet-I or mCherry; excitation dichroic mirror, FF458 Di-02 for FRET assay and FF409/493/573/652/759-Di01-25.8x37.8 for other purposes; emission filters, 445/40 for Calcofluor white, 483/32-25 for Turquoise2-GL, 520/28 for mNeonGreen, and 641/75 for mScarlet-I or mCherry (Semrock). The illumination light source was a Spectra X light engine (Lumencor). For snapshot imaging, cells were concentrated by centrifugation (3000 rpm, 1 min) and spotted on a glass slide (thickness, 0.9-1.2 mm; Matsunami). The specimen were sealed by a glass coverslip (thickness, 0.13-0.17 mm; Matsunami).

The ONIX Microfluidic platform (Merck) was used for long-term live-cell imaging. In a typical protocol, growing fission yeast cells were concentrated by centrifugation (3000 rpm, 1 min) and resuspended in 100 μL of the appropriate medium. The suspension was loaded into a trapping chamber (Y04T-04) by imposing pressure of 8 psi for 15 sec. During the imaging, the media was perfused by imposing pressure of 1 psi unless otherwise stated.

### Image analysis and data visualization

All the images were analyzed using Fiji/ImageJ^65^. The background was subtracted by the rolling-ball method (rolling ball radius, 50.0 pixels) adopted in Fiji. An optimized tracking plugin for Fiji, LIM Tracker^66^, enabled easy single-cell tracking. Regions of interest (ROIs) typically corresponded to the nucleus, as determined from CFP images. Each fluorescence intensity was defined as the mean value of ROIs. If the degree of aggregation was of interest, for example in the case of Pcn1, the coefficient of variation of the fluorescence intensity was calculated for that protein.

Manual image analysis was required for some experiments, typically for measuring cell length. For this, fission yeast cells were stained with calcofluor white (PhytoTech, C1933) after being washed twice with 1 mL DDW and centrifugation (3000 rpm, 1 min) when only the cell length was of interest. When the relation between CDK activity and cell length was of interest, this procedure was not used. Instead, DIC images were used to measure the length, because both the cross-talk of calcofluor white and the effect of the washing process on CDK activity deteriorated the accuracy of the analyses. In both cases, the length was manually determined using Fiji. “Length at division” and “Length at birth” were measured as the pole-to-pole length and the pole-to-septum length of cells forming a clear septum. To quantify the length during the vegetative cell cycle, we adopted the following two criteria. First, septated cells were considered as different cells, and the pole-to-septum length was defined as the length of each cell. Second, cells that proceeded in telophase but were not yet septated were considered as different cells sharing the same length, *i.e.*, the pole-to-pole length was measured for both divided nuclei.

Accumulated numerical data were visualized using Python with seaborn and matplotlib. The detailed procedures are described in the legend of each figure. The efficiency of FRET was quantified as the ratio of fluorescence intensity of YFP to that of CFP.

### Dose-response measurements

Liquid cultured cells in mid-log phase were dispensed to a fresh tube and further incubated with 1 μM 1-NM-PP1, an inhibitory analog of ATP that is specific of modified kinases^67^, for 3 h at 32°C. The cells were collected by centrifugation (3000 rpm, 30 sec) followed by washing with various concentrations of 1-NM-PP1 ranging from 10 nM to 10 μM. The washing process was repeated three times. The cell pellet was then suspended into the 20 μL each washing solution. The cell suspension was quickly spotted on a glass slide and observed under the microscope as described above. The images were taken within 15 min from the beginning of the washing process. Images were analyzed as follows. First, CFP images were binarized by Otsu’s method in Fiji. Second, the regions of the nucleus were detected by the “analyze particle” method in Fiji. The collected ROIs were overlaid on DIC images to exclude inadequate regions.

The obtained data were fitted with the following function using the Python library, scipy.optimize.curve_fit:

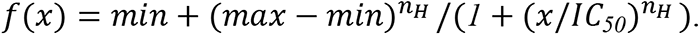

We gave the following initial values for the optimization:

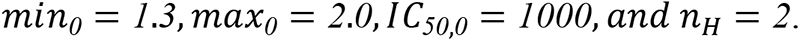

### Characterization of cell cycle events during the G2 and M phases

In Figure 3, we defined the following 7 events during the G2 and M phase from a nuclear division: #1, Cyclin degradation; #2, CDK deactivation; #3, Cell growth halt; #4, CDK peak; #5, Spindle emergence; #6, Cyclin peak; #7, CDK change-point. The events #1, #2, #3, #4, #5, and #7 were measured from the same imaging data obtained every 2 min. The event #6 (spindle emergence) was characterized using an independent experiment. The timing of these events was determined according to the following criteria.

1. Cyclin degradation: The time point when the fluorescence intensity of cyclin decreased most steeply. Time series data were adjusted by subtracting their previous values, and the time point that showed the minimum value was detected.
2. CDK deactivation: The time when the FRET/CFP ratio of Eevee-spCDK decreased most steeply. Time series data were adjusted by subtracting their previous values, and the time point that showed the minimum value was detected.
3. Cell growth halt: The time point when cell length reached a plateau during the G2 and M phases. The pole-to-pole cell length was manually measured using the DIC images. The obtained time series data were fitted with two linear functions, and the time at which the two lines intersect was determined.
4. CDK peak: The time point when the FRET/CFP ratio reaches its maximum over one cell cycle (from the first nuclear division to the next nuclear division).
5. Spindle emergence: The time point when a visible punctum was observed. Fluorescence time-lapse images of endogenous tubulin fused with mScarlet-I were taken every 1 min, and the time point when spindle tubulin emerged as a punctum was monitored manually.
6. Cyclin peak: The time point when the fluorescence intensity of Cdc13-mScarlet-I reaches its maximum over one cell cycle (from the first nuclear division to the next nuclear division).
7. CDK change-point: The time point when the steep activation of CDK started. The time series data of the FRET/CFP ratio of Eevee-spCDK were fitted with two linear functions, and the time at which the two lines intersect was determined.

### Western blotting

Protein extract was prepared based on previously reported methods^49, 68^. In brief, fission yeast cells were cultured at 32°C, and about 2.5 ∼ 5.0 × 10^7^ cells were collected by centrifugation. For cell lysis, the collected cells were suspended in 1 mL of ice-cold 20% TCA, and then washed with 1 mL of 1 M Tris-HCl (pH 7.5). The washed cells were suspended in 200 μL of 2 × SDS sample buffer (25 mM Tris-HCl (pH 6.8), 24% glycerol, 4% SDS, 0.008% BromoPhenol Blue, and 10% 2-mercaptoethanol) and incubated at 95°C for 10 min. Subsequently, the samples were transferred to a tube containing 7.0-mm zirconia beads (Bio Medical Science), disrupted using a Cell Destroyer PS2000 (Bio Medical Science), incubated at 95°C for 10 min, and centrifuged (16,000 g, 10 min, 4°C). The supernatants were loaded on 5∼20% polyacrylamide gels (Nacalai Tesque, cat #13063-74). Proteins were separated by SDS-PAGE and transferred to polyvinylidene difluoride membranes (Merck Millipore, cat #IPFL00010). After blocking with Intercept TBS Blocking Buffer (LI-COR, cat #927-60001), the membranes were probed with primary antibodies against FLAG-tag (dilution 1:1000, mouse; Sigma, A1804), Wee1 (dilution 1:1000, rabbit; Abcam, ab233540), and α-tubulin (dilution 1:1000, mouse; Sigma-Aldrich, T5168), followed by IRDye800CW donkey anti-mouse IgG secondary antibody (dilution 1:5000, LI-COR, 926-32212) and IRDye680RD goat anti-rabbit IgG secondary antibody (dilution 1:5000, LI-COR, 926-68071). Proteins were detected by an Odyssey infrared scanner (LI-COR).

### Measurement of the growth rate of fission yeast

Fission yeast cells were pre-cultured at 32℃ up to an optical density at 600 nm (OD600) of 1.0, followed by dilution to 1:20. A Compact Rocking Incubator Biophotorecorder TVS062CA (Advantec) was used for culture growth (32℃, 70 rpm) and OD660 measurement. Growth curves were fitted by the logistic function (*x = K / (1 + (K/x0 -1)e^-rt^)*), and doubling time (*ln2/ r*) was calculated.

## Acknowledgements

We thank all members of the Aoki Laboratory for their helpful discussions and assistance. We would also like to thank Paul M. Nurse and Nitin Kapadia for helpful feedback on the manuscript. Some fission yeast strains were provided by the National Bio-Resource Project (NBRP), Japan. K.A. was supported by JSPS KAKENHI Grants (nos. 18H02444, 19H05798, and 22H02625). Y.G. was supported by JSPS KAKENHI Grants (nos.19K16050 and 22K15110) and a Sumitomo Research grant. H.S. was supported by JSPS KAKENHI Grants (nos. 21J01354 and 22K15115) and the Dr. Yoshifumi Jigami Memorial Fund, The Society of Yeast Scientists. DC was supported by the Agence Nationale de la Recherche (PRC eVOLve, ANR-18-CD13-0009) and the Région Nouvelle Aquitaine (program CHESS, grant agreements 15963520 and 15964420).

## Competing Interests

No competing interests declared.

## Author contributions

Y.G. and K.A. designed the research. H.S. and Y.G. performed the experiments. D.C. assisted the experiments. H.S. and Y.G. analyzed the data. Y.K. constructed and analyzed the mathematical model. H.S., Y.G., and K.A. drafted the manuscript. All authors approved the final manuscript.

## Notes

### Competing Interest Statement

The authors have declared no competing interest.

