## Supplemental Information for "Live-cell imaging provides direct evidence for a threshold in CDK activity at the G2/M transition"

fission yeast, cyclin-dependent kinase, FRET, imaging

**Figure S1.**

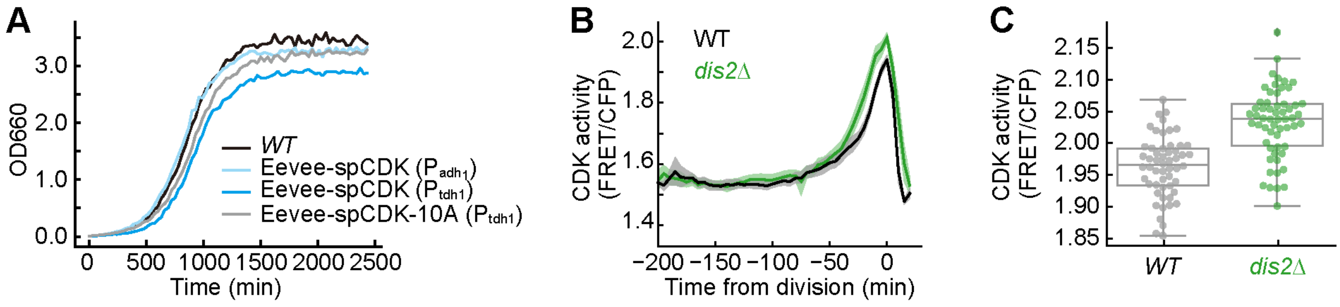


**Figure S1. Validation of Eevee-spCDK.**
(A) Growth curves of WT cells (black line), cells expressing Eevee-spCDK under the *adh1* promoter (light blue line), Eevee-spCDK under the *tdh1* promoter (dark blue line), and the non-phosphorylatable mutant Eevee-spCDK-10A under the *tdh1* promoter (gray line).
(B) Time courses of the FRET/CFP ratio in WT (black) and *dis2*-deleted mutant cells (*dis2Δ*, green). Shaded areas indicate the 95% confidence interval. Time 0 is defined as the timing of peak CDK activity.
(C) Peak values of the FRET/CFP ratio for WT (gray) and *dis2Δ* (green) cells. Each dot represents a single cell. Boxplots show the quartiles of the accumulated data with whiskers denoting the minimum and maximum except for the outliers detected as 1.5 times the interquartile range. n = 54 cells for WT; n = 58 cells for *dis2Δ*.

**Figure S2.**

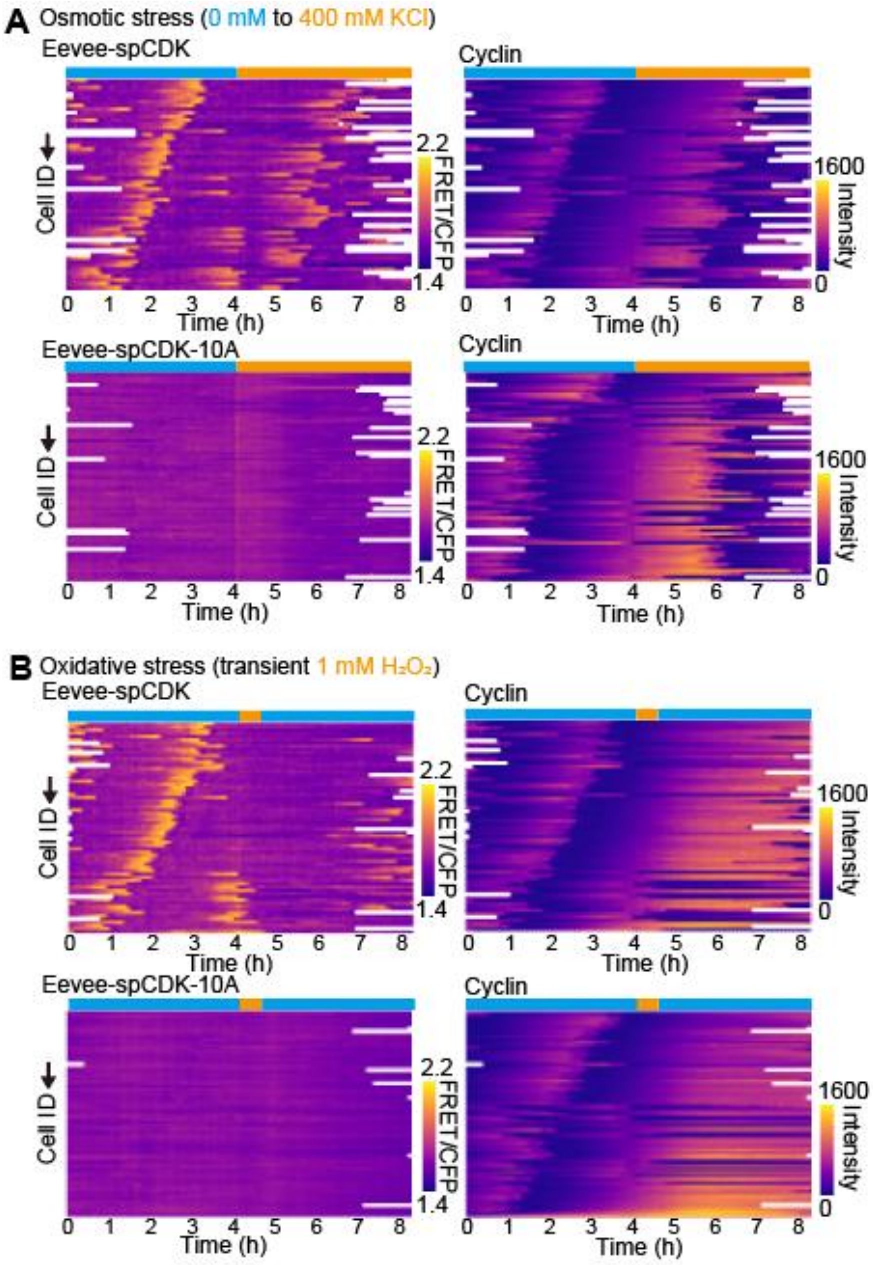


**Figure S2. Response of CDK activity to osmotic or oxidative stress.**
(A) Response of the FRET/CFP ratio of Eevee-spCDK or Eevee-spCDK-10A (left column) and cyclin level (right column) to an osmotic shock imposed by 400 mM KCl. Quantified time course data were aligned to the time point when the osmotic shock was applied (time 4). The data were sorted by the cyclin levels measured 30 min before the osmotic shock.
(B) Response of the FRET/CFP ratio of Eevee-spCDK or Eevee-spCDK-10A (left column) and the cyclin level (right column) to an oxidative shock imposed by transient (30 min) exposure to 1 mM $H_2O_2$. Quantified time course data were aligned by the time point when the oxidative shock was applied (time 0). The data were sorted by cyclin levels measured 30 min before the stress.

**Figure S3.**

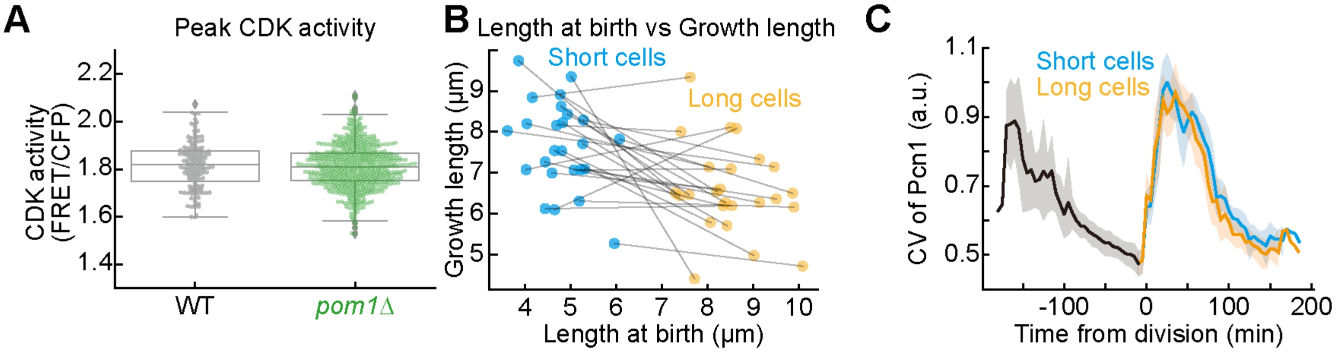


**Figure S3. Characterization of *pom1Δ* cells.**

(A) Peak CDK activity in WT (gray dots) and *pom1*-deleted mutant cells (*pom1Δ*, green dots). Each dot represents a single cell. Boxplots show the quartiles of the accumulated data with whiskers denoting the minimum and maximum except for the outliers detected as 1.5 times the interquartile range. n = 143 cells for WT; n = 560 cells for *pom1Δ*.

(B) Relation between length at birth and growth length in short (blue dots) and long (orange dots) *pom1Δ* mutant cells. Black lines connect corresponding sister cells. n = 25 cells.

(C) Comparison of S phase between the short (blue) and long (orange) *pom1Δ* mutant cells. Coefficient of variation (CV) of Pcn1-mScarlet-I was calculated (see Materials and Methods) and used as an S phase marker. Time 0 is defined by the timing of nuclear division. Lines show the mean CV values of Pcn1-mScalet-I in short (blue) and long (orange) *pom1Δ* mutant cells. The black line indicates their mother. Shaded areas indicate the 95% confidence interval. n = 9 cells. a.u.: arbitrary unit.

**Figure S4.**

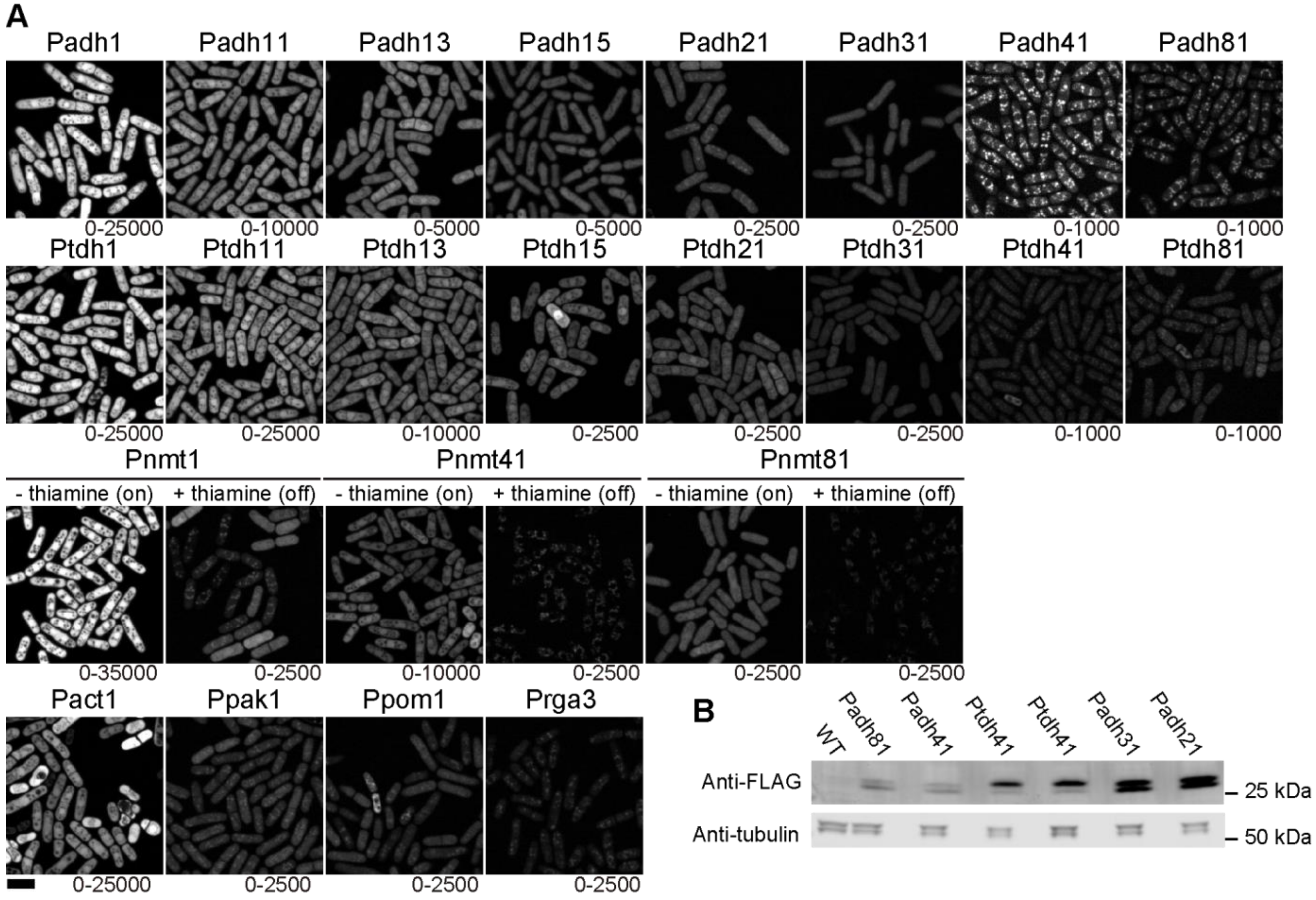


**Figure S4. Quantification of the strength of various promoters.**
(A) Representative fluorescence images for cells expressing mNeonGreen driven by the indicated promoters. For display purpose, the maximum and minimum values are modified for each image as shown at the bottom right of each image. Scale bar, 10 µm.
(B) Evaluation of the strength of the indicated promoters by Western blot. Cells expressing mNeonGreen-3xFLAG were lysed as described in the Materials and Methods section. The cell lysates were analyzed by Western blot with anti-FLAG (upper) and anti-tubulin (lower) antibodies.

**Figure S5.**

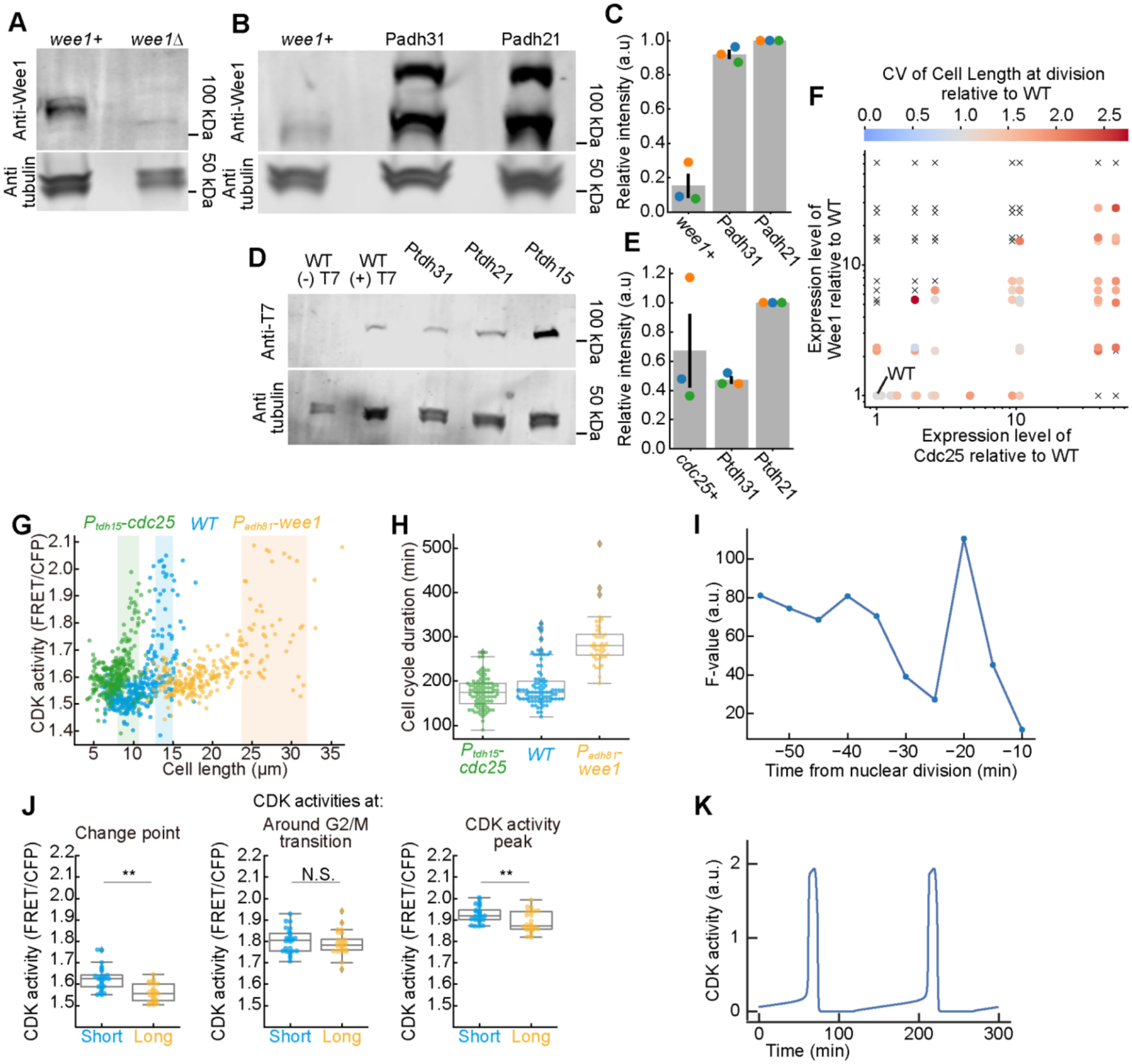


**Figure S5. Effects of the overexpressing of *cdc25* and/or *wee1* as well as deleting *pom1* on cell length, cell cycle duration, and CDK activity.**

(A) Validation of anti-Wee1 antibody. Cell lysates from WT (*wee1+*) and *wee1Δ* cells were analyzed by Western blot with anti-Wee1 (upper) and anti-tubulin (lower) antibodies.

(B and C) Quantification of Wee1 expression. (B) Cell lysates from WT (*wee1+*) and cells expressing Wee1-mScarlet-I driven by the indicated promoters were analyzed as in panel A. Of note, the presence of 2 bands in the lysates of the cells expressing Wee1-mScarlet-I suggest partial degradation of Wee1-mScarlet-I including the potential cleavage of mScarlet-I. Both bands were taken into account for the quantification. (C) Quantification of the protein amounts from Western blot as in B. The band intensities were normalized by the intensity derived from *Padh21*-driven Wee1-mScarlet-I (sum of two

bands), and the relative intensities were plotted. Each dot represents the value determined from an independent experiment. Error bars denote the standard error. n = 3.
(D and E) Quantification of Cdc25 expression. (D) Cell lysates from parental WT cells ((-) T7), or cells expressing Cdc25-mScarlet-I-T7 driven by the *cdc25* endogenous promoter ((+) T7) or the indicated promoters were analyzed by Western blot with anti-T7 (upper) or anti-tubulin (lower) antibodies. (E) Quantification of the protein amounts from Western blots as in D. The band intensities are normalized by *Ptdh21*-driven Cdc25-mScarlet-I-T7. The relative intensities are plotted. Each dot represents the value determined from an independent experiment. Error bars denote the standard error. n = 3.
(F) Coefficient of variation (CV) of cell length at division relative to WT. The expression level of each protein was obtained from the Western blots (see above). Each dot or cross mark corresponds to a strain harboring a specific combination of promoters for overexpressing *wee1* and *cdc25* at different levels. Cross marks represent combinations that lead to lethality. The relative CV values of cell length to WT in each strain are color-coded as shown at the top. n > 20 cells for each strain.
(G) Relation between cell length and CDK activity in WT cells (blue), cells overexpressing *cdc25* (green) or *wee1* (orange) as quantified from snapshot images. Each dot represents the data from a single cell. Pale color bands represent the ranges of cell length at division for the corresponding strains (the width of each band indicates the standard deviation). n = 353 for WT cells, n = 920 for *cdc25*-overexpressing cells, and n = 640 for *wee1*-overexpressing cells.
(H) Comparison of cell cycle duration between WT cells (blue), *cdc25*-overexpressing cells (green), and *wee1*-overexpressing cells (orange). Each dot represents the duration from a nuclear division to the next for each cell. Boxplots show the quartiles of the accumulated data with whiskers denoting the minimum and maximum except for the outliers detected as 1.5 times the interquartile range. n = 85 for WT cells, n = 89 for *cdc25*-overexpressing cells, and n = 44 for *wee1*-overexpressing cells.
(I) Variance of CDK activity between the strains (*cdc25*-overexpressing cells, *wee1*-overexpressing cells and WT) relative to that within the strains (F-value). F-values were quantified at various time points before nuclear division.
(J) CDK activity at the change-point (left), cyclin peak (middle), and CDK activity peak (right) in short (blue dots) and long (orange dots) *pom1Δ* mutant cells. Boxplots show the quartiles of the accumulated data with whiskers denoting the minimum and maximum except for the outliers detected as 1.5 times the interquartile range. n = 25 cells.
(K) Numerical simulation of the dynamics of CDK activity during the cell cycle. The value of CDK activity at each timepoint was calculated based on the ordinary differential equation as proposed in previous literature (see main text for the references).

**Figure S6.**

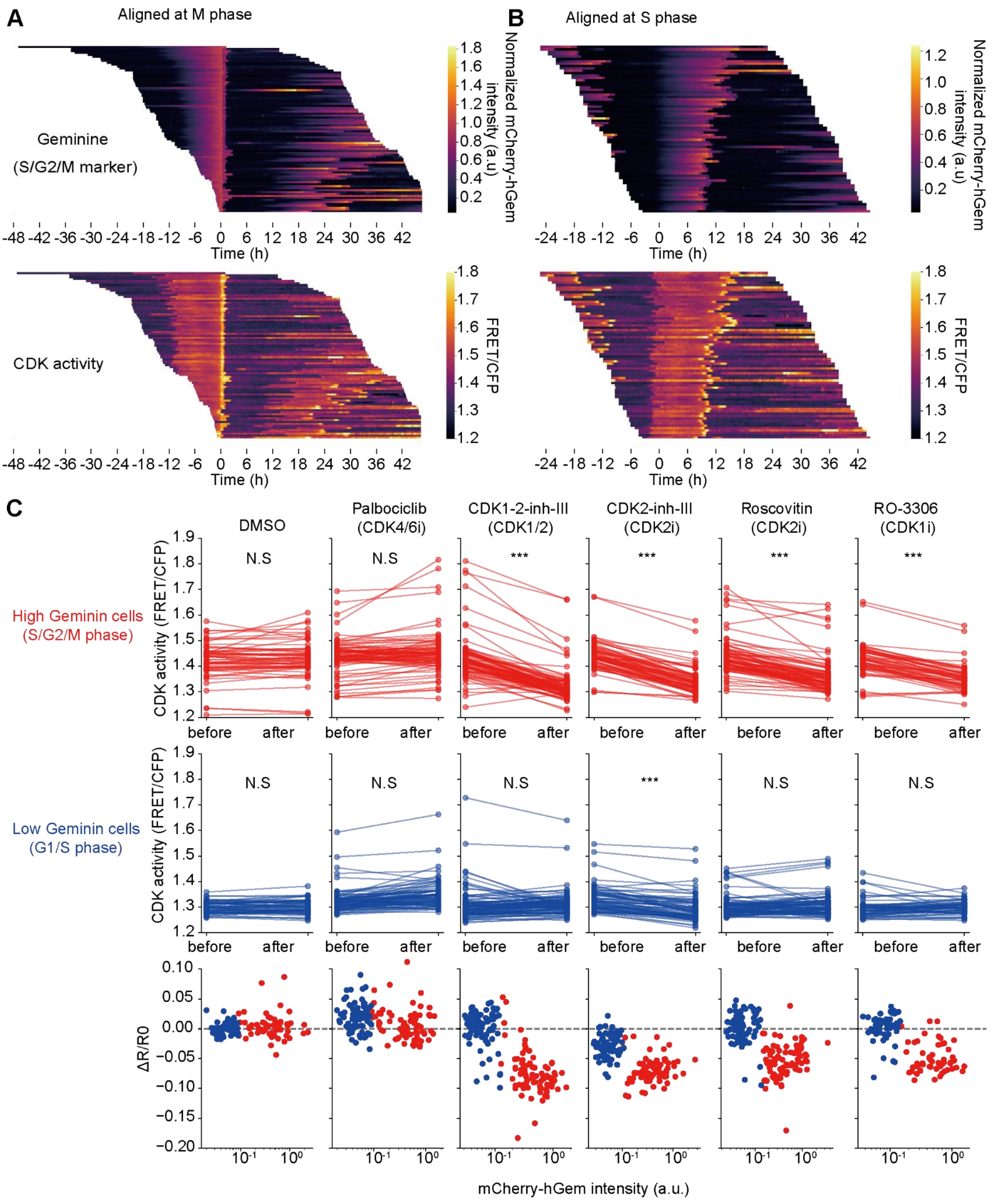


**Figure S6. Characterization of Eevee-spCDK as a CDK sensor in mammalian cells.**

(A and B) Heatmaps of normalized mCherry-Geminin intensity (upper) and FRET/CFP ratio of Eevee-spCDK (lower) aligned at M phase (A) or at S phase onset (B) in HeLa cells. HeLa cells expressing Eevee-spCDK and mCherry-Geminin were imaged for two days with a wide-field epifluorescence microscope. The timings of mitotic exit and S phase onset were assigned to the time of the sudden drop in mCherry-Geminin intensity and the start of Geminin accumulation, respectively.

(C) Effects of the indicated CDK inhibitors on the FRET/CFP ratio of Eevee-spCDK in HeLa cells. HeLa cells expressing Eevee-spCDK and mCherry-Geminin were imaged with a wide-field epifluorescence microscope. The cells were treated with DMSO or the indicated CDK inhibitors for 30 min. For all inhibitors, a concentration of 10 μM was used. The FRET/CFP ratios were obtained before and after CDK inhibitor treatment. Furthermore, the cells were further divided into two groups: high-Geminin cells (S/G2/M cells, red, top) and low-Geminin cells (G1 cells, blue, middle). Each dot represents the FRET/CFP ratio. Lines link the ratios measured for the same cell before and after treatments. One-sided paired t-tests were performed (N.S: not significant, $p > 0.05$; ***: $p < 0.001$). Changes in the FRET/CFP ratio before and after treatment ($\Delta R/R0$) are plotted against normalized Geminin intensity (bottom). If an inhibitor decreases the FRET/CFP ratio, $\Delta R/R0$ should be below zero. Red dots indicate high-Geminin cells (S/G2/M cells), and blue dots indicate low-Geminin cells (G1 cells).

**Movie S1. CDK activity and cyclin expression level during the cell cycle.** The FRET/CFP ratio of Eevee-spCDK in the fission yeast *S. pombe* is represented in the intensity-modulated display mode with pseudo-colors, the scale of which are shown at the lower side of the video. The images were taken every 5 min. The timestamp shows time in hours: minutes. Scale bar, 10 μm.

**Movie S2. CDK activity and heterogeneity of Pcn1 distribution in the nucleus for visualizing S phase.** The FRET/CFP ratio of Eevee-spCDK in the fission yeast *S. pombe* is represented in the intensity-modulated display mode with pseudo-colors, the scale of which are shown at the lower side of the video. The heterogeneous distribution of Pcn1-mScarlet-I indicates S phase. The images were taken every 5 min. The timestamp shows time in hours: minutes. Scale bar, 10 μm.

**Movie S3. Response of CDK activity and cyclin expression level to osmotic shock.** The osmotic shock was imposed by treating cells with 400 mM KCl at time = 50 min in the video. The FRET/CFP ratio of Eevee-spCDK in the fission yeast *S. pombe* is represented in the intensity-modulated display mode with pseudo-colors, the scale of which are shown at the lower side of the video. The images were taken every 5 min. The timestamp shows time in hours: minutes. Scale bar, 10 μm.

**Movie S4. Response of CDK activity and cyclin expression level to oxidative shock.** The oxidative shock was imposed by a transient (30 min) exposure to 1 mM $H_2O_2$ at time = 50 min in the video**.** The FRET/CFP ratio of Eevee-spCDK in the fission yeast *S. pombe* is represented in the intensity-modulated display mode with pseudo-colors, the scale of which are shown at the lower side of the video. The images were taken every 5 min. The timestamp shows time in hours: minutes. Scale bar, 10 μm.

**Movie S5. CDK activity and mCherry-hGem expression level during the cell cycle.** The FRET/CFP ratio of Eevee-spCDK in HeLa cells is represented in the intensity-modulated display mode with pseudo-colors, the scale of which are shown at the lower side of the video. The images were taken every 10 min. The timestamp shows time in hours: minutes. Scale bar, 20 μm.

**Table S1. Plasmids constructed and used in this study.**

| Plasmid name | Description | Source | Benchling Link |
|---|---|---|---|
| pSKI(ver.2)-NAT-2L-A1-M | pSKI-v2 | this study | https://benchling.com/s/seq-ueKfUe6rue5pfddos6LH?m=slm-WHzkMYJAIFF38S0d2zyo |
| pSKI(ver.2)-NAT-2L-A11-M | pSKI-v2 | this study | https://benchling.com/s/seq-X384jWMZBAdgYF0DB4Mj?m=slm-1A7V5m0n50lssRb0Cn4t |
| pSKI(ver.2)-NAT-2L-A13-M | pSKI-v2 | this study | https://benchling.com/s/seq-tmaH7AeD4jMTZv7PJhxk?m=slm-YbOSeiIcURlqx4xLhapy |
| pSKI(ver.2)-NAT-2L-A15-M | pSKI-v2 | this study | https://benchling.com/s/seq-E9nqx40UjqqB7u8wtfNZ?m=slm-yC61Bo3JI72MTx8QTsRI |
| pSKI(ver.2)-NAT-2L-A21-M | pSKI-v2 | this study | https://benchling.com/s/seq-gXpE8sdH4pJn0uPqAga1?m=slm-LAJ2G9snMVE6g9re7H5O |
| pSKI(ver.2)-NAT-2L-A31-M | pSKI-v2 | this study | https://benchling.com/s/seq-tkRFpfmSfDEyr0D7amae?m=slm-MUZshf0mmTFicjwGvl4f |
| pSKI(ver.2)-NAT-2L-A41-M | pSKI-v2 | this study | https://benchling.com/s/seq-MML1zXApeD7Yi1R2OqQM?m=slm-QqQUDqf7AHc7GAicukPA |
| pSKI(ver.2)-NAT-2L-A81-M | pSKI-v2 | this study | https://benchling.com/s/seq-6vtKcJ9TEQCz3BzGE0uk?m=slm-VK6npU9etFrrlDsbwXjJ |
| pSKI(ver.2)-NAT-2L-T1-M | pSKI-v2 | this study | https://benchling.com/s/seq-kKpCKm5iENrVJJcw6m3U?m=slm-72JdVqlp8IAVk6LWiGG8 |
| pSKI(ver.2)-NAT-2L-T11-M | pSKI-v2 | this study | https://benchling.com/s/seq-4n3C1hEtFPUfdRr3K89X?m=slm-RxTsbHOAuVU5zgxhrbbz |
| pSKI(ver.2)-NAT-2L-T13-M | pSKI-v2 | this study | https://benchling.com/s/seq-9Cf4InguLrI4lPWOjbBS?m=slm-INuG3fnXUd4CpsF9A5Gp |
| pSKI(ver.2)-NAT-2L-T15-M | pSKI-v2 | this study | https://benchling.com/s/seq-O7g3iksSI6RythSsPe7Q?m=slm-2rTQfgTL9n4KGAjRaeux |
| pSKI(ver.2)-NAT-2L-T21-M | pSKI-v2 | this study | https://benchling.com/s/seq-GnZwxKqUxphwx6JEe6WV?m=slm-iI808jE5cexWkhelMAWk |
| pSKI(ver.2)-NAT-2L-T31-M | pSKI-v2 | this study | https://benchling.com/s/seq-vPD413SNKNh6MgUZE1uB?m=slm-DnofGujGeAWyJkWdeDmh |

| | | | |
|---|---|---|---|
| pSKI(ver.2)-NAT-2L-T41-M | pSKI-v2 | this study | https://benchling.com/s/seq-L1uiuV5LfhbHkdci2Sxo?m=slm-oKb7iZPvGw4yhzCqygvW |
| pSKI(ver.2)-NAT-2L-T81-M | pSKI-v2 | this study | https://benchling.com/s/seq-JGO074gfN24DYRmTm6q4?m=slm-ub316fZeDrKFIRh1jA0E |
| pSKI(ver.2)-NAT-2L-AC1-M | pSKI-v2 | this study | https://benchling.com/s/seq-ci8aka3L3LgLRkxfk9dl?m=slm-p5gOGDU3nmgmDsnSH5UF |
| pSKI(ver.2)-NAT-2L-PA1-M | pSKI-v2 | this study | https://benchling.com/s/seq-rPIizZ2wDoy6nX1u7VHt?m=slm-V750F8mzxTEJeCShhBAG |
| pSKI(ver.2)-NAT-2L-R3-M | pSKI-v2 | this study | https://benchling.com/s/seq-8BYqxDPaQdmVjYTBVKvZ?m=slm-mB2oBFHkz7m6iGCQ1Mn4 |
| pSKI(ver.2)-NAT-2L-PO1-M | pSKI-v2 | this study | https://benchling.com/s/seq-DXpcEOLp6mwWUmXPscjx?m=slm-sonb18WIjxxMzbqgUm1D |
| pSKI(ver.2)-NAT-2L-N1-M | pSKI-v2 | this study | https://benchling.com/s/seq-kKKEHbr8aLKsfEJMAThW?m=slm-nBNHeUyy5Is1xIz0IKhM |
| pSKI(ver.2)-NAT-2L-N41-M | pSKI-v2 | this study | https://benchling.com/s/seq-9eaR9B69KuUaVSXFL9KO?m=slm-6UYVlESzQwQ4u6w6GtuD |
| pSKI(ver.2)-NAT-2L-N81-M | pSKI-v2 | this study | https://benchling.com/s/seq-ySLik4OWCZBW5Bd9dsAj?m=slm-Q1cxM1fAA52sa3Ea9PDf |
| pSKI(ver.2)-KAN-3R-A1-M | pSKI-v2 | this study | https://benchling.com/s/seq-l6CBOtBvK3dHIBzgMinS?m=slm-cNAlkwqjaYhhM18AvvL7 |
| pSKI(ver.2)-KAN-3R-A11-M | pSKI-v2 | this study | https://benchling.com/s/seq-hQZZ2NalhEfQhXqI08kH?m=slm-Z7ifJ0oRa5f4VWhaIXiN |
| pSKI(ver.2)-KAN-3R-A13-M | pSKI-v2 | this study | https://benchling.com/s/seq-YZImX7ogPoLnKYAAatMC?m=slm-sxWKnViQ1vanVSnUbAHD |
| pSKI(ver.2)-KAN-3R-A15-M | pSKI-v2 | this study | https://benchling.com/s/seq-5WP4AjyDUCPdcnmRlUb6?m=slm-hLLYbc6kv9kSJdW0Ndk4 |
| pSKI(ver.2)-KAN-3R-A21-M | pSKI-v2 | this study | https://benchling.com/s/seq-Cn9Dr0Hu3PCA5AltvYrp?m=slm-PfwFXZ2hY6cqiYPqsiHy |
| pSKI(ver.2)-KAN-3R-A31-M | pSKI-v2 | this study | https://benchling.com/s/seq-Qeq6QpBOfzxVYEURyvhm?m=slm-xpkUlvaeyZIyPzopP8ae |

| | | | |
|---|---|---|---|
| pSKI(ver.2)-KAN-3R-A41-M | pSKI-v2 | this study | https://benchling.com/s/seq-2yadjE8URc0NG9gb7Ue5?m=slm-B6O7XWlpapteSjrPb4Hh |
| pSKI(ver.2)-KAN-3R-A81-M | pSKI-v2 | this study | https://benchling.com/s/seq-80IJsyMaICYwYNOCLG7Z?m=slm-zXKfPNAboOQsWqbZ6n6u |
| pSKI(ver.2)-KAN-3R-T1-M | pSKI-v2 | this study | https://benchling.com/s/seq-FDhGaMldAePlBRBXPTny?m=slm-OWrQKI2RhAHmk3My0ITP |
| pSKI(ver.2)-KAN-3R-T11-M | pSKI-v2 | this study | https://benchling.com/s/seq-KsbCk2osKH1pVuSXw0JV?m=slm-AMEUFWdUpsjMtNhJuvIa |
| pSKI(ver.2)-KAN-3R-T13-M | pSKI-v2 | this study | https://benchling.com/s/seq-fWaI49GPxYFPl9bXT1iR?m=slm-7mP00ThzrI55uefXO7Tj |
| pSKI(ver.2)-KAN-3R-T15-M | pSKI-v2 | this study | https://benchling.com/s/seq-bxdqh4EGXdnjdYKE0Gcw?m=slm-amg5GTAo8btqGYmgNjzN |
| pSKI(ver.2)-KAN-3R-T21-M | pSKI-v2 | this study | https://benchling.com/s/seq-0ir8TGZgOJU4a4aKYrNV?m=slm-IUbYHmBgowJku86y80o5 |
| pSKI(ver.2)-KAN-3R-T31-M | pSKI-v2 | this study | https://benchling.com/s/seq-B8DNuLog32UjWPG3asd4?m=slm-KXQWJ9q1BP87ufkNvl9W |
| pSKI(ver.2)-KAN-3R-T41-M | pSKI-v2 | this study | https://benchling.com/s/seq-yDfhpMHq3UDPBCRtr2MH?m=slm-JSdqy4PPtVctkZQ8KjaD |
| pSKI(ver.2)-KAN-3R-T81-M | pSKI-v2 | this study | https://benchling.com/s/seq-bKOmBXjzSXc5FCNFBYHi?m=slm-1NeBT4FpaduGDcVCmZcc |
| pSKI(ver.2)-KAN-3R-AC1-M | pSKI-v2 | this study | https://benchling.com/s/seq-R5Qz15IVp3H7lR6YOp8z?m=slm-ZFgU0ge756DanPcDfq7f |
| pSKI(ver.2)-KAN-3R-PA1-M | pSKI-v2 | this study | https://benchling.com/s/seq-oIDMuB5ua7ERzNPCLkJP?m=slm-JxaqWwlhM1mb5sV5cQ5O |
| pSKI(ver.2)-KAN-3R-R3-M | pSKI-v2 | this study | https://benchling.com/s/seq-pyYDZ5pRspWTOUEMT74A?m=slm-kpmuC9LNMEFeVyJL56TB |
| pSKI(ver.2)-KAN-3R-PO1-M | pSKI-v2 | this study | https://benchling.com/s/seq-wieeoPp0TwRlxD5rq8me?m=slm-fqKUi7TlvYawfrI809vR |
| pSKI(ver.2)-KAN-3R-N1-M | pSKI-v2 | this study | https://benchling.com/s/seq-TAYTEGSyAiOY0HeRIAc3?m=slm-3dkGxF80flhXLSygoehw |

| | | | |
|---|---|---|---|
| pSKI(ver.2)-KAN-3R-N41-M | pSKI-v2 | this study | https://benchling.com/s/seq-KP3x0D3gwkmWVuRbaimi?m=slm-qXtwDGhs9QQlyQDSwx02 |
| pSKI(ver.2)-KAN-3R-N81-M | pSKI-v2 | this study | https://benchling.com/s/seq-d1eGAeBNg6lXbZyXIpYx?m=slm-gmjju4vjfRXwzNp6XJ2e |
| pSKI(ver.2)-BSD-1L-A1-M | pSKI-v2 | this study | https://benchling.com/s/seq-zAK4GISbkHQQbAL42XU4?m=slm-bxns2nOmxqeTVZQe3XI2 |
| pSKI(ver.2)-BSD-1L-T1-M | pSKI-v2 | this study | https://benchling.com/s/seq-UHJtcqRaD5PVyxlF5611?m=slm-F24RUzQaT2oqG9dg4aq3 |
| pSKI(ver.2)-NAT-2L-A1-M-mNeonGreen | | this study | https://benchling.com/s/seq-H2HF4jWahPswMyH6SAsX?m=slm-9QldpZY8vBMVTjWelgj8 |
| pSKI(ver.2)-NAT-2L-A11-M-mNeonGreen | | this study | https://benchling.com/s/seq-Rd1UgufoHC2TNPaokfs1?m=slm-F9VAZU2w1JKdhnhxLB7Z |
| pSKI(ver.2)-NAT-2L-A13-M-mNeonGreen | | this study | https://benchling.com/s/seq-rEkwASflhzWz55sEGCqj?m=slm-TqI81qXcG873hFT9M8l9 |
| pSKI(ver.2)-NAT-2L-A15-M-mNeonGreen | | this study | https://benchling.com/s/seq-Z9FyYetpnCULxpGrgCws?m=slm-S7badT0G5DjOqDKvbYXE |
| pSKI(ver.2)-NAT-2L-A21-M-mNeonGreen | | this study | https://benchling.com/s/seq-v8AcsPRtAo64f43zm6a3?m=slm-GGcgUC0tizYU2ZpxLzmO |
| pSKI(ver.2)-NAT-2L-A31-M-mNeonGreen | | this study | https://benchling.com/s/seq-IfBJkrqK2V8MDE54QulN?m=slm-tZjjfwwZpR0jnmw7tt3j |
| pSKI(ver.2)-NAT-2L-A41-M-mNeonGreen | | this study | https://benchling.com/s/seq-mvloeca0Q0e03okIcAdF?m=slm-U7OaWG8YIK3vQWPjYLkB |
| pSKI(ver.2)-NAT-2L-A81-M-mNeonGreen | | this study | https://benchling.com/s/seq-oQ2psy665DVKyXkia642?m=slm-pZZ6QSxoqBgDhfMBKmd4 |
| pSKI(ver.2)-NAT-2L-T1-M-mNeonGreen | | this study | https://benchling.com/s/seq-mPBFzfzbHik3tCi0VsAY?m=slm-VRv28mw2xn3UkpZo8yoj |
| pSKI(ver.2)-NAT-2L-T11-M-mNeonGreen | | this study | https://benchling.com/s/seq-txQEPh1yjUTb0QQPWDbG?m=slm-F3odB7b4x9NVLWNZSJBD |
| pSKI(ver.2)-NAT-2L-T13-M-mNeonGreen | | this study | https://benchling.com/s/seq-GUqvGIPeyeKT8EUBjUeL?m=slm-7s2XCJ7Hp1xnYuxAzcVj |

| | | | |
|---|---|---|---|
| pSKI(ver.2)-NAT-2L-T15-M-mNeonGreen | | this study | https://benchling.com/s/seq-0KudjnWLS93IbiiGaf2Z?m=slm-oC4iXJZ65hrXnjNgdBZ5 |
| pSKI(ver.2)-NAT-2L-T21-M-mNeonGreen | | this study | https://benchling.com/s/seq-gyplbD4UHwKfWzUJXR8y?m=slm-lVG8rPqkMql707HkIphr |
| pSKI(ver.2)-NAT-2L-T31-M-mNeonGreen | | this study | https://benchling.com/s/seq-nXhVA4uj4CP8efzgTrQW?m=slm-QFfaVx36giTqVCf4MQEi |
| pSKI(ver.2)-NAT-2L-T41-M-mNeonGreen | | this study | https://benchling.com/s/seq-wb0ySafQPRSp3qIOFppC?m=slm-RPERKKcMm7op3ZsILgrf |
| pSKI(ver.2)-NAT-2L-T81-M-mNeonGreen | | this study | https://benchling.com/s/seq-uG2jlrutN3YASv4giZEX?m=slm-n3j6jrft1fx4FuLD0h06 |
| pSKI(ver.2)-NAT-2L-AC1-M-mNeonGreen | | this study | https://benchling.com/s/seq-yzh6OTsUI8fugNJyILZ7?m=slm-HKdc24KV5xsdtSFvRRzy |
| pSKI(ver.2)-NAT-2L-PA1-M-mNeonGreen | | this study | https://benchling.com/s/seq-9wzn2twFYCZkn0XYIHSV?m=slm-smHkt0SUHZH2DoPchPnx |
| pSKI(ver.2)-NAT-2L-R3-M-mNeonGreen | | this study | https://benchling.com/s/seq-3AD93X5NQpkck0DezLQl?m=slm-2dNyGW2wUv9ZAJiGzzFc |
| pSKI(ver.2)-NAT-2L-PO1-M-mNeonGreen | | this study | https://benchling.com/s/seq-l4C1JfZpaAbAXFVEzYrt?m=slm-hkdgr7ctYLBQSl6oYSA3 |
| pSKI(ver.2)-NAT-2L-N1-M-mNeonGreen | | this study | https://benchling.com/s/seq-FMxFvmoozaCYVbPwt1oC?m=slm-z2wBr7SMv5gu91GijooX |
| pSKI(ver.2)-NAT-2L-N41-M-mNeonGreen | | this study | https://benchling.com/s/seq-mGO2drXioyVWTsiNlBbK?m=slm-mHeZSG8SlX3elQPtQmEo |
| pSKI(ver.2)-NAT-2L-N81-M-mNeonGreen | | this study | https://benchling.com/s/seq-ThrD9Orw0RmdYXHlZfnr?m=slm-qhQAJjwK4tkWrSUOxBtp |
| pSKI(ver.2)-NAT-2L-A21-M-mNeonGreen-3FLAG | | this study | https://benchling.com/s/seq-R1VT7F69Brr2oxstOuyR?m=slm-ReEoTMtuXYECFuSoGnjH |
| pSKI(ver.2)-NAT-2L-A31-M-mNeonGreen-3FLAG | | this study | https://benchling.com/s/seq-sM2vegZv0OWXJd6BnGwa?m=slm-iZmPm7AMGSd7OOjFFXK4 |
| pSKI(ver.2)-NAT-2L-A41-M-mNeonGreen-3FLAG | | this study | https://benchling.com/s/seq-bnu7tQseLQPBjlOCh78m?m=slm-QtduUb3uBXfqzfrnfdCV |

| | | | |
|---|---|---|---|
| pSKI(ver.2)-NAT-2L-A81-M-mNeonGreen-3FLAG | | this study | https://benchling.com/s/seq-fRosnUyGHocHuMRy8xap?m=slm-p99hQCGVmbk0RiVCqLce |
| pSKI(ver.2)-NAT-2L-T41-M-mNeonGreen-3FLAG | | this study | https://benchling.com/s/seq-HiTnZiRiwpCEQoNGxPyF?m=slm-HRw6Qx14OtAaUOlFLaJF |
| pSKI(ver.2)-NAT-2L-T81-M-mNeonGreen-3FLAG | | this study | https://benchling.com/s/seq-lWiG93NOJs6A7bsIil3W?m=slm-DqR7TMbMMKmyVgFuezHT |
| pSKI(ver.2)-NAT-2L-A1-M-cdc25-mScarlet-I | | this study | https://benchling.com/s/seq-SzrgyemV2ywmg7ELaTgX?m=slm-SVTprvLSz6TfmSH884pt |
| pSKI(ver.2)-NAT-2L-A11-M-cdc25-mScarlet-I | | this study | https://benchling.com/s/seq-DJ3wExRtt1AQCw1gumEK?m=slm-iGPAbceBAi1BlrpIjGGH |
| pSKI(ver.2)-NAT-2L-A13-M-cdc25-mScarlet-I | | this study | https://benchling.com/s/seq-UPJ6ZPr9TaIf534aEjxR?m=slm-YhfCO4EXSYhyq98nUh74 |
| pSKI(ver.2)-NAT-2L-A15-M-cdc25-mScarlet-I | | this study | https://benchling.com/s/seq-qa9pTw9LHHnXdOCbQBpM?m=slm-hWq1qMOP3WDWYAza1hpi |
| pSKI(ver.2)-NAT-2L-A21-M-cdc25-mScarlet-I | | this study | https://benchling.com/s/seq-qyfw5c1THsmioJQ7GObK?m=slm-mnwgPOn9jWluYs1kXVI0 |
| pSKI(ver.2)-NAT-2L-A31-M-cdc25-mScarlet-I | | this study | https://benchling.com/s/seq-SecSWG77H47C5TvEUOEb?m=slm-nVoyhwZxdHuoAxEGei2X |
| pSKI(ver.2)-NAT-2L-A41-M-cdc25-mScarlet-I | | this study | https://benchling.com/s/seq-d09s5BEoxOjhtQtZI67K?m=slm-94NJfQEHmRvVCKyJeXlc |
| pSKI(ver.2)-NAT-2L-A81-M-cdc25-mScarlet-I | | this study | https://benchling.com/s/seq-V7VR5KtJqrSG04KSKnbb?m=slm-VHTeQ4W199Ns3D1qWZVI |
| pSKI(ver.2)-NAT-2L-T1-M-cdc25-mScarlet-I | | this study | https://benchling.com/s/seq-1DQZlX4FDfMwmKVXkmml?m=slm-tO5aUsHja0Nl2fTEvExF |
| pSKI(ver.2)-NAT-2L-T11-M-cdc25-mScarlet-I | | this study | https://benchling.com/s/seq-OzsiJ2ruQ1jLAg5E6RAK?m=slm-aSPc9q1qIEO5GYPpC4Mf |
| pSKI(ver.2)-NAT-2L-T13-M-cdc25-mScarlet-I | | this study | https://benchling.com/s/seq-8AofsJ6cYIubPZvQJc9u?m=slm-9eSW3pTN52XKo4U1GtGu |
| pSKI(ver.2)-NAT-2L-T15-M-cdc25-mScarlet-I | | this study | https://benchling.com/s/seq-kJ07CNtfIsGhKStFQDx2?m=slm-iJ8T1OkqpyYVFRMtjeWX |

| | | | |
|---|---|---|---|
| pSKI(ver.2)-NAT-2L-T21-M-cdc25-mScarlet-I | | this study | https://benchling.com/s/seq-ssrtS1CauVlUo44UmS5l?m=slm-f4o4tK8YBdr1shPKOQw9 |
| pSKI(ver.2)-NAT-2L-T31-M-cdc25-mScarlet-I | | this study | https://benchling.com/s/seq-9z5ZKnQ1OQOH80YFpLVJ?m=slm-rFFZ6UnwGLtWJYN5eKU1 |
| pSKI(ver.2)-NAT-2L-T41-M-cdc25-mScarlet-I | | this study | https://benchling.com/s/seq-PhPaJjbCk7qEZWp5zQAM?m=slm-XufgSxkIJvPW4zJUHOBf |
| pSKI(ver.2)-NAT-2L-T81-M-cdc25-mScarlet-I | | this study | https://benchling.com/s/seq-BxwiipIV99n8mMo5UFDS?m=slm-jkPJIi9KlkiTcxNq1pms |
| pSKI-KAN-3R-A11-M-wee1-mScarlet-I | | this study | https://benchling.com/s/seq-aZF3dyqTuoEtzxliCGAf?m=slm-a1KvbdxADdmKUXT1daw8 |
| pSKI(ver.2)-KAN-3R-A11-M-wee1-mScarlet-I | | this study | https://benchling.com/s/seq-Kji7RJe8ZNecBjs8aCSr?m=slm-ZYUJDnE92wVkg6zr0W6g |
| pSKI(ver.2)-KAN-3R-A13-M-wee1-mScarlet-I | | this study | https://benchling.com/s/seq-Fja1hn7sFV6ghN7cLu4z?m=slm-L8rYPKY7tP8huxwd7v37 |
| pSKI(ver.2)-KAN-3R-A15-M-wee1-mScarlet-I | | this study | https://benchling.com/s/seq-cuLrDBQkknMCHEDqZZL1?m=slm-wfsvsTkhu5veIpzdz4Fg |
| pSKI(ver.2)-KAN-3R-A21-M-wee1-mScarlet-I | | this study | https://benchling.com/s/seq-ehqbSPPqlbnK2QLJKqSa?m=slm-GgKnSAbi9gAX1tIZnzof |
| pSKI(ver.2)-KAN-3R-A31-M-wee1-mScarlet-I | | this study | https://benchling.com/s/seq-wLeAEsWgqZKDhr3CzyrN?m=slm-6x16N4MuBzrzaRZMZ4Xq |
| pSKI(ver.2)-KAN-3R-A41-M-wee1-mScarlet-I | | this study | https://benchling.com/s/seq-tPpusS6Dxl2NcG7sFjjq?m=slm-6LA1b4IM4yioAZem4xcK |
| pSKI(ver.2)-KAN-3R-A81-M-wee1-mScarlet-I | | this study | https://benchling.com/s/seq-hIn0tJvxY9x1WOlzNctY?m=slm-2EG1f5l5VMmOVMIYVDut |
| pSKI(ver.2)-KAN-3R-T1-M-wee1-mScarlet-I | | this study | https://benchling.com/s/seq-zjnWiyTy9JGUeb2acbPR?m=slm-2o3tJBB8WqBY5krhTBy8 |
| pSKI(ver.2)-KAN-3R-T11-M-wee1-mScarlet-I | | this study | https://benchling.com/s/seq-AJB0kTrMHwWCROkGkwJR?m=slm-ohAnurax9MaHXBGm1KBc |
| pSKI(ver.2)-KAN-3R-T13-M-wee1-mScarlet-I | | this study | https://benchling.com/s/seq-69ckGW5xxym1YwLKdYs4?m=slm-y3lnmtx2FkJBcD7a2HTn |

| | | | |
|---|---|---|---|
| pSKI(ver.2)-KAN-3R-T15-M-wee1-mScarlet-I | | this study | https://benchling.com/s/seq-Y8KL5cYIkgZSwE3NRy86?m=slm-gzP4W5ccLi5T1I4OBrb9 |
| pSKI(ver.2)-KAN-3R-T21-M-wee1-mScarlet-I | | this study | https://benchling.com/s/seq-V2xHbl16emoUsBZ8u1Fa?m=slm-VvXzw7wNSljj8yp8oxTr |
| pSKI(ver.2)-KAN-3R-T31-M-wee1-mScarlet-I | | this study | https://benchling.com/s/seq-DHbGFA78ht5mh4W3Vzvk?m=slm-zYBr7nIKMeGQvmYpwWs9 |
| pSKI(ver.2)-KAN-3R-T41-M-wee1-mScarlet-I | | this study | https://benchling.com/s/seq-tpJVhctQdMdo57jfpS7Z?m=slm-aJC0zzd6F43GxnG2722Y |
| pSKI(ver.2)-KAN-3R-T81-M-wee1-mScarlet-I | | this study | https://benchling.com/s/seq-1ifGkvpQTRLALryXu1WR?m=slm-lqcAMVAfCZt6Ldx1sTCH |
| pMNATZA1-EV-spCDK-nls | | this study | https://benchling.com/s/seq-YEccBRx9WcOm8HFC64iZ |
| pSKI-NAT-1L-A1-EV-spCDK-nls | | this study | https://benchling.com/s/seq-Ydf6KVJnWbOnazf9suWr?m=slm-tgKaTXef0RygdHVn7z4v |
| pSKI(ver.2)-NAT-2L-T1-M-EV-spCDK-nls | | this study | https://benchling.com/s/seq-xXgtsoJuTH1svGyr8ITU?m=slm-LFtGbN6OlxkbfYubq3hQ |
| pSKI-BSD-1L-A1-EV-spCDK-nls | | this study | https://benchling.com/s/seq-CAN0RcnSa01eR4VfVB8Q?m=slm-4zABPwdGZwlMQXaQdJYG |
| pSKI-BSD-2L-A1-EV-spCDK-nls | | this study | https://benchling.com/s/seq-sS3g1f6Em8MlE1kCYhZZ?m=slm-2TMAJyllm56L9mKV74WS |
| pSKI(ver.2)-NAT-2L-T1-M-EV-spCDK-10A-nls | | this study | https://benchling.com/s/seq-RY7RCzJxiovNt5cttnTX?m=slm-Mes0j1IELxXwZ6lQFUKV |
| pSKI-KAN-1L-A1-synCut3-mScarlet-I | | this study | https://benchling.com/s/seq-BKGo9GJRUEXYCHnP59J5 |
| pFA6a-mScarlet-I-nat | | this study | https://benchling.com/s/seq-Ukk7iYZEizrRCCswniDh |
| pFA6a-mScarlet-I-bsd | | this study | https://benchling.com/s/seq-eEGW5XmaWzlDBbQQ1xPM |
| pFA6a-mScarlet-I-kan | | this study | https://benchling.com/s/seq-f0m0oRjnOFr2HHo51waH |
| pFA6a-3FLAG-bsd | | this study | https://benchling.com/s/seq-C29p4RoTIzSpfUUAgnyd |
| pPBbsr2-EV-spCDK-nls | for human cell line | this study | https://benchling.com/s/seq-fqxUFt6oCznkbghFAC5N |
| pCSIIhyg-H2B-iRFP-P2A-mCherry-hGem | for human cell line | this study | https://benchling.com/s/seq-KF7uAXN7AhH28cnctu3l |

| | | | |
|---|---|---|---|
| pMNATZ1-EV-screening-01 (cut3-1 NLS) | | this study | https://benchling.com/s/seq-0qFCP0UqAnTo9F6f6Zpy?m=slm-MC7eDLYQpNiqmPGN55sI |
| pMNATZ1-EV-screening-02 (cut3-2 NLS) | | this study | https://benchling.com/s/seq-3lN8D4StDKrJqo87AMTr?m=slm-purkf5oxYqIaW5Uw6DLz |
| pMNATZ1-EV-screening-03 (cut3-3 NLS) | | this study | https://benchling.com/s/seq-n4Trn4huwT4HSEnPJnOE?m=slm-vNS9NU7fWLXD9HJis3uD |
| pMNATZ1-EV-screening-04 (alp14-1 NLS) | | this study | https://benchling.com/s/seq-Xw1cTi0sJJgGacZ9olDW?m=slm-qO3fsxppMjLD5DeUjZcE |
| pMNATZ1-EV-screening-05 (alp14-2 NLS) | | this study | https://benchling.com/s/seq-UiEEXRlOfooyH78nvrPS?m=slm-Yfb5hJbHqutTOf35dIQp |
| pMNATZ1-EV-screening-06 (ask1 NLS) | | this study | https://benchling.com/s/seq-yUMWDEBxyVv29NwUPIUw?m=slm-S7AVrurDwgkSx7j36s7y |
| pMNATZ1-EV-screening-07 (rad2 NLS) | | this study | https://benchling.com/s/seq-ILiph6dMiw8TzUpagEBV?m=slm-DRXxIpGlPDI6RPRDGWXl |
| pMNATZ1-EV-screening-08 (orc1 NLS) | | this study | https://benchling.com/s/seq-VJxHonXZ53jcAJmeNswy?m=slm-NX13ZFpP1o90gVUEZHnD |
| pMNATZ1-EV-screening-09 (orc2 NLS) | | this study | https://benchling.com/s/seq-tNOpt2J7rZOlmff3JlsM?m=slm-yfV3rophvzj6iMVBweUE |
| pMNATZ1-EV-screening-10 (cut3N NLS) | | this study | https://benchling.com/s/seq-mIfqTDy3JRRr8x5nNwLt?m=slm-KDbBbeg5p6UNTDNR2MKJ |
| pMNATZ1-EV-screening-11 (drc1-1 NLS) | | this study | https://benchling.com/s/seq-naZiekIt4kW31DMPtRUo?m=slm-BNaOPgB7mUmio52asfNt |
| pMNATZ1-EV-screening-12 (drc1-2 NLS) | | this study | https://benchling.com/s/seq-xreesbvGEJkWOUFKjG0m?m=slm-OwWskPLTgCnb7CZQDPbr |
| pMNATZ1-EV-screening-13 (drc1-3 NLS) | | this study | https://benchling.com/s/seq-dyyZvoXtUZuUzPD4tBI7?m=slm-7um6Fnw9harZxGMhn8rT |

**Table S2. S. pombe strain genotypes used in this study.**

| Strain name | Genotype | Fig. | Source |
|---|---|---|---|
| L972 | h- | Fig. 4A, B, Fig. S4B, Fig.S5A, B, C, D | NBRP |
| L968 | h90 | Fig.S1A | NBRP |
| HS034 | h90 cdc13-mScarlet-I<<kan | - | this study, L968 |
| HS048 | h- 1L::Padh1-Eevee-spCDK-nls<<bsd | - | this study, L972 |
| DC240 | h+ leu1Δ::Pcdc13::cdc13-L-cdc2as::cdc13 3'UTR::ura4+ cdc2Δ::kanMX6 cdc13Δ::natMX6 cig1Δ::ura4+ cig2Δ::ura4+ puc1Δ::ura4+ ura4-D18 | - | Damien et al.,2010 |
| AN202 | h- cdc13-mScarlet-I<<bsd | - | this study, L972 |
| HS364 | h- cdc13-spmScarlet-I<<bsd pom1::nat | - | this study, AN202 |
| HS406 | h- dis2::nat | - | this study, L972 |
| YG1369 | h- cdc13-mScarlet-I<<bsd z::Padh1-Eevee-spCDK-candidate-nls-025(cut3-1 NLS)<<nat | Fig. 1C | this study, AN202 |
| YG1370 | h- cdc13-mScarlet-I<<bsd z::Padh1-Eevee-spCDK-candidate-nls-026(cut3-2 NLS)<<nat | Fig. 1C | this study, AN202 |
| YG1371 | h- cdc13-mScarlet-I<<bsd z::Padh1-Eevee-spCDK-candidate-nls-027(cut3-3 NLS)<<nat | Fig. 1C | this study, AN202 |
| YG1372 | h- cdc13-mScarlet-I<<bsd z::Padh1-Eevee-spCDK-candidate-nls-028(alp14-1 NLS)<<nat | Fig. 1C | this study, AN202 |
| YG1373 | h- cdc13-mScarlet-I<<bsd z::Padh1-Eevee-spCDK-candidate-nls-029(alp14-2 NLS)<<nat | Fig. 1C | this study, AN202 |
| YG1374 | h- cdc13-mScarlet-I<<bsd z::Padh1-Eevee-spCDK-candidate-nls-030(ask1 NLS)<<nat | Fig. 1C | this study, AN202 |
| YG1375 | h- cdc13-mScarlet-I<<bsd z::Padh1-Eevee-spCDK-candidate-nls-031(rad2 NLS)<<nat | Fig. 1C | this study, AN202 |
| YG1376 | h- cdc13-mScarlet-I<<bsd z::Padh1-Eevee-spCDK-candidate-nls-032(orc1 NLS)<<nat | Fig. 1C | this study, AN202 |
| YG1377 | h- cdc13-mScarlet-I<<bsd z::Padh1-Eevee-spCDK-candidate-nls-033(orc2 NLS)<<nat | Fig. 1C | this study, AN202 |
| YG1381 | h- cdc13-mScarlet-I<<bsd z::Padh1-Eevee-spCDK-candidate-nls-037(cut3N NLS)<<nat | Fig. 1C | this study, AN202 |
| HS351 | h- cdc13-mScarlet-I<<bsd Z::Padh1-Eevee-spCDK-candidate-nls-024 (Drc1-N NLS)<<nat | Fig. 1C | this study, AN202 |

| | | | |
|---|---|---|---|
| HS007 | h90 z::Padh1-Eevee-spCDK-nls<<nat cdc13-mScarlet-I<<kan | Fig. 1E, F, G, Fig2. C, D, E, H, I, J, K, Fig. 3A, B, Fig. S2A, B, Fig. S3A | this study HS001 |
| HS012 | h90 z::Pahd1-Eevee-spCDK-nls<<nat pcn1-mScarlet-I<<kan | Fig. 2F | this study HS001 |
| HS013 | h90 z::Padh1-Eevee-spCDK-nls<<nat 3R::Padh11-synCut3-mScarlet-I<<kan | Fig. 2G | this study HS001 |
| HS078 | h- 2L::Padh1-mNeonGreen<<nat | Fig. 5B, Fig. S4A | this study, L972 |
| HS079 | h- 2L::Padh11-mNeonGreen<<nat | Fig. 5B, Fig. S4A | this study, L972 |
| HS080 | h- 2L::Padh13-mNeonGreen<<nat | Fig. 5B, Fig. S4A | this study, L972 |
| HS081 | h- 2L::Padh15-mNeonGreen<<nat | Fig. 5B, Fig. S4A | this study, L972 |
| HS082 | h- 2L::Padh21-mNeonGreen<<nat | Fig. 5B, Fig. S4A | this study, L972 |
| HS083 | h- 2L::Padh31-mNeonGreen<<nat | Fig. 5B, Fig. S4A | this study, L972 |
| HS084 | h- 2L::Padh41-mNeonGreen<<nat | Fig. 5B, Fig. S4A | this study, L972 |
| HS085 | h- 2L::Padh81-mNeonGreen<<nat | Fig. 5B, Fig. S4A | this study, L972 |
| HS086 | h- 2L::Ptdh1-mNeonGreen<<nat | Fig. 5B, Fig. S4A | this study, L972 |
| HS087 | h- 2L::Ptdh11-mNeonGreen<<nat | Fig. 5B, Fig. S4A | this study, L972 |
| HS105 | h- 2L::Ptdh13-mNeonGreen<<nat | Fig. 5B, Fig. S4A | this study, L972 |
| HS088 | h- 2L::Ptdh15-mNeonGreen<<nat | Fig. 5B, Fig. S4A | this study, L972 |
| HS089 | h- 2L::Ptdh21-mNeonGreen<<nat | Fig. 5B, Fig. S4A | this study, L972 |
| HS090 | h- 2L::Ptdh31-mNeonGreen<<nat | Fig. 5B, Fig. S4A | this study, L972 |
| HS091 | h- 2L::Ptdh41-mNeonGreen<<nat | Fig. 5B, Fig. S4A | this study, L972 |
| HS092 | h- 2L::Ptdh81-mNeonGreen<<nat | Fig. 5B, Fig. S4A | this study, L972 |
| HS093 | h- 2L::Pact1-mNeonGreen<<nat | Fig. 5B, Fig. S4A | this study, L972 |
| HS094 | h- 2L::Ppak1-mNeonGreen<<nat | Fig. 5B, Fig. S4A | this study, L972 |
| HS095 | h- 2L::Prga3-mNeonGreen<<nat | Fig. 5B, Fig. S4A | this study, L972 |

| | | | |
|---|---|---|---|
| HS096 | h- 2L::Ppom1-mNeonGreen<<nat | Fig. 5B, Fig. S4A | this study, L972 |
| HS097 | h- 2L::Pnmt1-mNeonGreen<<nat | Fig. 5B, Fig. S4A | this study, L972 |
| HS098 | h- 2L::Pnmt41-mNeonGreen<<nat | Fig. 5B, Fig. S4A | this study, L972 |
| HS099 | h- 2L::Pnmt81-mNeonGreen<<nat | Fig. 5B, Fig. S4A | this study, L972 |
| HS231 | h- 2L::Padh21-mNeonGreen-3FLAG<<nat | Fig. 5C, FigS4B | this study, L972 |
| HS232 | h- 2L::Padh31-mNeonGreen-3FLAG<<nat | Fig. 5C, FigS4B | this study, L972 |
| HS233 | h- 2L::Padh41-mNeonGreen-3FLAG<<nat | Fig. 5C, FigS4B | this study, L972 |
| HS234 | h- 2L::Padh81-mNeonGreen-3FLAG<<nat | Fig. 5C, FigS4B | this study, L972 |
| HS237 | h- 2L::Ptdh41-mNeonGreen-3FLAG<<nat | Fig. 5C, FigS4B | this study, L972 |
| HS238 | h- 2L::Ptdh81-mNeonGreen-3FLAG<<nat | Fig. 5C, FigS4B | this study, L972 |
| HS049 | h90 2L::Padh11-cdc25-mScarlet-I<<nat | Fig. 5E, S5F | this study, L968 |
| HS050 | h90 2L::Padh13-cdc25-mScarlet-I<<nat | Fig. 5E, S5F | this study, L968 |
| HS051 | h90 2L::Padh15-cdc25-mScarlet-I<<nat | Fig. 5E, S5F | this study, L968 |
| HS052 | h90 2L::Padh21-cdc25-mScarlet-I<<nat | Fig. 5E, S5F | this study, L968 |
| HS053 | h90 2L::Padh31-cdc25-mScarlet-I<<nat | Fig. 5E, S5F | this study, L968 |
| HS054 | h90 2L::Padh41-cdc25-mScarlet-I<<nat | Fig. 5E, S5F | this study, L968 |
| HS055 | h90 2L::Padh81-cdc25-mScarlet-I<<nat | Fig. 5E, S5F | this study, L968 |
| HS106 | h90 2L::Ptdh13-cdc25-mScarlet-I<<nat | Fig. 5E, S5F | this study, L968 |
| HS107 | h90 2L::Ptdh15-cdc25-mScarlet-I<<nat | Fig. 5E, S5F | this study, L968 |
| HS108 | h90 2L::Ptdh21-cdc25-mScarlet-I<<nat | Fig. 5E, S5F | this study, L968 |
| HS109 | h90 2L::Ptdh31-cdc25-mScarlet-I<<nat | Fig. 5E, S5F | this study, L968 |
| HS110 | h90 2L::Ptdh41-cdc25-mScarlet-I<<nat | Fig. 5E, S5F | this study, L968 |

| | | | |
|---|---|---|---|
| HS111 | h90 2L::Ptdh81-cdc25-mScarlet-I<<nat | Fig. 5E, S5F | this study, L968 |
| HS112 | h90 3R::Padh41-wee1-mScarlet-I<<kan | Fig. 5E, S5F | this study, L968 |
| HS113 | h90 3R::Padh81-wee1-mScarlet-I<<kan | Fig. 5E, S5F | this study, L968 |
| HS140 | h90 2L::Ptdh13-cdc25-mScarlet-I<<nat 3R::Padh21-wee1-mScarlet-I<<kan | Fig. 5E, S5F | this study, HS106 |
| HS141 | h90 2L::Ptdh13-cdc25-mScarlet-I<<nat 3R::Padh31-wee1-mScarlet-I<<kan | Fig. 5E, S5F | this study, HS106 |
| HS142 | h90 2L::Ptdh13-cdc25-mScarlet-I<<nat 3R::Padh41-wee1-mScarlet-I<<kan | Fig. 5E, S5F | this study, HS106 |
| HS143 | h90 2L::Ptdh13-cdc25-mScarlet-I<<nat 3R::Padh81-wee1-mScarlet-I<<kan | Fig. 5E, S5F | this study, HS106 |
| HS168 | h90 2L::Ptdh13-cdc25-mScarlet-I<<nat 3R::Ptdh21-wee1-mScarlet-I<<kan | Fig. 5E, S5F | this study, HS106 |
| HS169 | h90 2L::Ptdh13-cdc25-mScarlet-I<<nat 3R::Ptdh31-wee1-mScarlet-I<<kan | Fig. 5E, S5F | this study, HS106 |
| HS146 | h90 2L::Ptdh13-cdc25-mScarlet-I<<nat 3R::Ptdh41-wee1-mScarlet-I<<kan | Fig. 5E, S5F | this study, HS106 |
| HS147 | h90 2L::Ptdh13-cdc25-mScarlet-I<<nat 3R::Ptdh81-wee1-mScarlet-I<<kan | Fig. 5E, S5F | this study, HS106 |
| HS148 | h90 2L::Ptdh15-cdc25-mScarlet-I<<nat 3R::Padh15-wee1-mScarlet-I<<kan | Fig. 5E, S5F | this study, HS107 |
| HS170 | h90 2L::Ptdh15-cdc25-mScarlet-I<<nat 3R::Padh21-wee1-mScarlet-I<<kan | Fig. 5E, S5F | this study, HS107 |
| HS171 | h90 2L::Ptdh15-cdc25-mScarlet-I<<nat 3R::Padh31-wee1-mScarlet-I<<kan | Fig. 5E, S5F | this study, HS107 |
| HS151 | h90 2L::Ptdh15-cdc25-mScarlet-I<<nat 3R::Padh41-wee1-mScarlet-I<<kan | Fig. 5E, S5F | this study, HS107 |
| HS152 | h90 2L::Ptdh15-cdc25-mScarlet-I<<nat 3R::Padh81-wee1-mScarlet-I<<kan | Fig. 5E, S5F | this study, HS107 |
| HS153 | h90 2L::Ptdh15-cdc25-mScarlet-I<<nat 3R::Ptdh31-wee1-mScarlet-I<<kan | Fig. 5E, S5F | this study, HS107 |
| HS154 | h90 2L::Ptdh15-cdc25-mScarlet-I<<nat 3R::Ptdh41-wee1-mScarlet-I<<kan | Fig. 5E, S5F | this study, HS107 |
| HS155 | h90 2L::Ptdh15-cdc25-mScarlet-I<<nat 3R::Ptdh81-wee1-mScarlet-I<<kan | Fig. 5E, S5F | this study, HS107 |
| HS172 | h90 2L::Ptdh21-cdc25-mScarlet-I<<nat 3R::Padh31-wee1-mScarlet-I<<kan | Fig. 5E, S5F | this study, HS108 |
| HS157 | h90 2L::Ptdh21-cdc25-mScarlet-I<<nat 3R::Padh41-wee1-mScarlet-I<<kan | Fig. 5E, S5F | this study, HS108 |
| HS158 | h90 2L::Ptdh21-cdc25-mScarlet-I<<nat 3R::Padh81-wee1-mScarlet-I<<kan | Fig. 5E, S5F | this study, HS108 |

| | | | |
|---|---|---|---|
| HS159 | h90 2L::Ptdh21-cdc25-mScarlet-I<<nat 3R::Ptdh31-wee1-mScarlet-I<<kan | Fig. 5E, S5F | this study, HS108 |
| HS160 | h90 2L::Ptdh21-cdc25-mScarlet-I<<nat 3R::Ptdh41-wee1-mScarlet-I<<kan | Fig. 5E, S5F | this study, HS108 |
| HS161 | h90 2L::Ptdh21-cdc25-mScarlet-I<<nat 3R::Ptdh81-wee1-mScarlet-I<<kan | Fig. 5E, S5F | this study, HS108 |
| HS162 | h90 2L::Ptdh31-cdc25-mScarlet-I<<nat 3R::Padh1-wee1-mScarlet-I<<kan | Fig. 5E, S5F | this study, HS109 |
| HS163 | h90 2L::Ptdh31-cdc25-mScarlet-I<<nat 3R::Padh11-wee1-mScarlet-I<<kan | Fig. 5E, S5F | this study, HS109 |
| HS164 | h90 2L::Ptdh31-cdc25-mScarlet-I<<nat 3R::Padh41-wee1-mScarlet-I<<kan | Fig. 5E, S5F | this study, HS109 |
| HS165 | h90 2L::Ptdh31-cdc25-mScarlet-I<<nat 3R::Padh81-wee1-mScarlet-I<<kan | Fig. 5E, S5F | this study, HS109 |
| HS166 | h90 2L::Ptdh31-cdc25-mScarlet-I<<nat 3R::Ptdh41-wee1-mScarlet-I<<kan | Fig. 5E, S5F | this study, HS109 |
| HS167 | h90 2L::Ptdh31-cdc25-mScarlet-I<<nat 3R::Ptdh81-wee1-mScarlet-I<<kan | Fig. 5E, S5F | this study, HS109 |
| HS262 | h90 2L::Ptdh13-cdc25-mScarlet-I<<nat 3R::Padh11-wee1-mScarlet-I<<kan | Fig. 5E, S5F | this study, L968 |
| HS263 | h90 2L::Ptdh13-cdc25-mScarlet-I<<nat 3R::Padh41-wee1-mScarlet-I<<kan | Fig. 5E, S5F | this study, L968 |
| HS264 | h90 2L::Ptdh13-cdc25-mScarlet-I<<nat 3R::Padh81-wee1-mScarlet-I<<kan | Fig. 5E, S5F | this study, L968 |
| HS265 | h90 2L::Ptdh13-cdc25-mScarlet-I<<nat 3R::Ptdh41-wee1-mScarlet-I<<kan | Fig. 5E, S5F | this study, L968 |
| HS266 | h90 2L::Ptdh11-cdc25-mScarlet-I<<nat 3R::Padh15-wee1-mScarlet-I<<kan | Fig. 5E, S5F | this study, L968 |
| HS267 | h90 2L::Ptdh11-cdc25-mScarlet-I<<nat 3R::Padh21-wee1-mScarlet-I<<kan | Fig. 5E, S5F | this study, L968 |
| HS268 | h90 2L::Ptdh11-cdc25-mScarlet-I<<nat 3R::Padh31-wee1-mScarlet-I<<kan | Fig. 5E, S5F | this study, L968 |
| HS269 | h90 2L::Ptdh11-cdc25-mScarlet-I<<nat 3R::Padh41-wee1-mScarlet-I<<kan | Fig. 5E, S5F | this study, L968 |
| HS270 | h90 2L::Ptdh11-cdc25-mScarlet-I<<nat 3R::Padh81-wee1-mScarlet-I<<kan | Fig. 5E, S5F | this study, L968 |
| HS271 | h90 2L::Ptdh11-cdc25-mScarlet-I<<nat 3R::Ptdh21-wee1-mScarlet-I<<kan | Fig. 5E, S5F | this study, L968 |
| HS272 | h90 2L::Ptdh11-cdc25-mScarlet-I<<nat 3R::Ptdh31-wee1-mScarlet-I<<kan | Fig. 5E, S5F | this study, L968 |
| HS273 | h90 2L::Ptdh11-cdc25-mScarlet-I<<nat 3R::Ptdh41-wee1-mScarlet-I<<kan | Fig. 5E, S5F | this study, L968 |
| HS274 | h90 2L::Ptdh11-cdc25-mScarlet-I<<nat 3R::Ptdh81-wee1-mScarlet-I<<kan | Fig. 5E, S5F | this study, L968 |

| | | | |
|---|---|---|---|
| HS296 | h90 2L::Ptdh1-cdc25-mScarlet-I<<nat 3R::Padh41-wee1-mScarlet-I<<kan | Fig. 5E, S5F | this study, L968 |
| HS297 | h90 2L::Ptdh1-cdc25-mScarlet-I<<nat 3R::Padh31-wee1-mScarlet-I<<kan | Fig. 5E, S5F | this study, L968 |
| HS298 | h90 2L::Ptdh1-cdc25-mScarlet-I<<nat 3R::Padh21-wee1-mScarlet-I<<kan | Fig. 5E, S5F | this study, L968 |
| HS299 | h90 2L::Ptdh1-cdc25-mScarlet-I<<nat 3R::Padh15-wee1-mScarlet-I<<kan | Fig. 5E, S5F | this study, L968 |
| HS300 | h90 2L::Ptdh1-cdc25-mScarlet-I<<nat 3R::Ptdh41-wee1-mScarlet-I<<kan | Fig. 5E, S5F | this study, L968 |
| HS301 | h90 2L::Ptdh1-cdc25-mScarlet-I<<nat 3R::Ptdh31-wee1-mScarlet-I<<kan | Fig. 5E, S5F | this study, L968 |
| HS302 | h90 2L::Ptdh1-cdc25-mScarlet-I<<nat 3R::Ptdh21-wee1-mScarlet-I<<kan | Fig. 5E, S5F | this study, L968 |
| HS303 | h90 2L::Ptdh1-cdc25-mScarlet-I<<nat 3R::Ptdh13-wee1-mScarlet-I<<kan | Fig. 5E, S5F | this study, L968 |
| HS189 | h90 1L::Padh1-Eevee-spCDK-nls<<bsd 2L::Ptdh15-cdc25-mScarlet-I<<nat | Fig. 5F, G, Fig. S5G, H | this study, HS107 |
| HS115 | h- 1L::Padh1-Eevee-spCDK-nls<<bsd 3R::Padh81-wee1-mScarlet-I<<kan | Fig. 5F, G, Fig. S5G, H | this study, HS048 |
| HS001 | h90 z::Padh1-Eevee-spCDK-nls<<nat | Fig. 5F, G, Fig. S1A, B, C, Fig. S5G, H | this study, L968 |
| HS360 | h90 z::Padh1-Eevee-spCDK-nls<<nat cdc13-mScarlet-I<<kan pom1::hyg | Fig. 4C, D, E, F, G, Fig. S3A, B | this study, HS007 |
| HS352 | h90 pom1::kan | Fig. 4A, B | this study, L968 |
| MY1757 | h- leu1 ura4 wee1::ura4 | Fig. S4A | NBRP |
| HS038 | h90 cdc13-mScarlet-I<<kan 2L::Ptdh1-Eevee-spCDK-nls-10A<<nat | Fig. 2H, J, Fig. S2A, B | this study, HS034 |
| HS019 | h+ leu1Δ::Pcdc13::cdc13-L-cdc2as::cdc13 3'UTR::ura4+ cdc2Δ::kanMX6 cdc13Δ::natMX6 cig1Δ::ura4+ cig2Δ::ura4+ puc1Δ::ura4+ ura4-D18 1L::Padh1-Eevee-spCDK-nls<<bsd | Fig. 2A, B | this study, DC240 |
| HS035 | h90 2L::Ptdh1-Eevee-spCDK-nls<<nat | Fig. S1A | this study, L968 |
| HS037 | h90 2L::Ptdh1-Eevee-spCDK-nls-10A<<nat | Fig. S1A | this study, L968 |
| HS349 | h90 2L::Ptdh1-cdc25-mScarlet-I<<nat 3R::Padh21-wee1-mScarlet-I<<kan 1L::Padh1-Eevee-spCDK-nls<<bsd | Fig. S4A, B, C | this study, HS298 |
| HS410 | h- dis2::nat 2L::Padh1(ver. 2)-CDKEV-024<<bsd | Fig. S1B, C | this study, HS406 |
| HS365 | h- cdc13-spmScarlet-I<<bsd pom1::nat pcn1-spmNeonGreen<<kan | Fig. S3C | this study, HS364 |

| | | | |
|---|---|---|---|
| HS306 | h90 z::Padh1-CDKEV024<<nat C::Padh15-mCherry-atb2<<hyg | Fig. 3A, B | this study, HS001 |
